# A systematic evaluation of yeast sample preparation protocols for spectral identifications, proteome coverage and post-isolation modifications

**DOI:** 10.1101/2022.01.17.476533

**Authors:** Maxime den Ridder, Ewout Knibbe, Wiebeke van den Brandeler, Pascale Daran-Lapujade, Martin Pabst

## Abstract

The importance of obtaining comprehensive and accurate information from cellular proteomics experiments asks for a systematic investigation of sample preparation protocols, particularly when working with unicellular organisms with strong cell walls, such as found in the model organism and cell factory *S. cerevisiae*. Sample preparation protocols may bias towards specific protein fractions or challenge the analysis of native protein modifications due to reagent-induced artefacts. Here, we performed a systematic comparison of sample preparation protocols using a matrix of different conditions commonly applied in whole cell lysate proteomics. The different protocols were evaluated for their overall fraction of identified spectra, proteome and amino acid sequence coverage, GO-term distribution and number of peptide modifications, by employing a combination of database and unrestricted modification search approaches. The best proteome and amino acid sequence coverage was achieved by using Urea combined with filter-aided or in-solution digestion protocols, where the overall outcomes were strongly influenced by the employed quenching procedure. Most importantly, the use of moderate incubation temperatures and times, circumvented excessive formation of modification artefacts. Extensive reagent-induced peptide modifications, however, were observed when using solvents such as acetone or additives such as formic acid. Moreover, several filter material-related modifications were observed when employing the filter-aided procedures. Ultimately, the best protocols enabled the identification of approximately 65–70% of all acquired fragmentation spectra, where additional *de novo* sequencing suggests that unidentified spectra were largely of too low spectral quality to provide confident spectrum matches. This study demonstrates the large impact of different sample preparation procedures on the proteomic analysis outcome, where the collected protocols and large sets of associated mass spectrometric raw data provide a resource to evaluate and design new protocols and guide the analysis of (native) peptide modifications in the model eukaryote yeast.

## INTRODUCTION

Despite recent advancements in mass spectrometric instrumentation, a large fraction of fragmentation spectra (MS/MS) from bottom-up shotgun proteomics experiments usually remain unidentified [1, 2]. Amongst the many possible reasons, a decreased identification rate can result from the presence of non-peptidic contaminants (particularly surfactants), increased spectral complexity due to co-fragmentation of multiple precursor ions, low-quality spectra due to poor and incomplete peptide fragmentation, unexpected peptide sequence variants or co/post-translational modifications that are not covered by the database, as well as deficiencies of the database search scoring schemes used to match the spectra. The latter led to the development of iterative approaches or to the combined use of orthogonal database search algorithms [3–7]. However, a significant fraction of unassigned MS/MS spectra may also be a consequence of incomplete or nonspecific proteolytic cleavage, or result from unintended peptide modifications introduced during sample preparation when using highly reactive chemicals. In fact, sample preparation can be one of the most significant contributors to data variation and poor comparability between proteomics experiments [8]. Hence, selecting the most suitable sample preparation protocol is essential for enabling a deep proteome coverage and for confidently identifying and quantifying (novel) peptide modifications. Moreover, the (native) *in vivo* state of the proteins should be preserved as good as possible. Therefore, sample preparation protocols usually include quenching of the cellular metabolism subsequently after sampling. This is in particular of importance, when investigating reversible post-translational modifications, which may be rapidly cleaved by the many hydrolytic enzymes present in every cell [9]. However, different quenching strategies, such as using ethanol or trichloroacetic acid (TCA) [10, 11], have been shown to differently impact the outcome of proteomics experiments [11]. Following protein extraction, proteins are solubilised and commonly denatured (e.g., with Urea), disulphide bonds are reduced by e.g., with Dithiothreitol (DTT) or Tris(2-carboxyethyl)phosphine (TCEP), alkylated using reagents such as iodoacetamide (IAA) or acrylamide (AA) to avoid reoxidation of the sulfhydryl groups [12, 13] and finally proteolytically cleaved (typically with trypsin) to create protein fragments that are sufficiently small for efficient measurement, but that retain sufficient unique sequence information for the subsequent protein identification step. Chemicals, or their employed concentrations, are often not compatible with the following steps in the protocol and, therefore, require removal by methods such as protein precipitation e.g., with acetone or Trichloroacetic acid (TCA), or filter-aided approaches [14–20]. The type and pH of the buffer used during the final proteolytic digestion may ultimately also impact the amino acid cleavage site specificity [21]. Finally, non-peptidic compounds and non-volatile salts are removed before analysis by the aid of solid-phase extraction (SPE), using different types of reverse-phase or mixed-mode resins [22]. However, combinations of different sample preparation procedures are expected to differently bias the proteomic analysis outcome. The selected protocol, therefore, may influence the proteome fraction or native modifications that can be detected and quantified significantly [23]. In addition, when investigating the biological significance of peptide modifications, the observed modification needs to be traced back to its co/post-translational or sample processing origin. However, without appropriate controls, this is a highly challenging procedure. For example, formylation can occur as a natural histone modification [24], but may also be introduced during sample preparation when using formic acid-containing buffers, such as those used to increase the solubility of hydrophobic peptides [25, 26]. Similarly for carbamylation, this modification has been associated with severe renal and cardiovascular disorders; however, it may also originate from high molarity Urea-containing buffers used during protein denaturation [27]. Alkylation of cysteine residues has become standard practice in proteomics experiments, which however due to over- or off-target alkylation reactions frequently introduces unintended peptide modifications, such as double alkylation artefacts, which for the case of IAA (114.04 Da) mimics the C-terminal glycine residue of ubiquitin [12, 13, 28, 29]. Nevertheless, recent studies suggest that disulphide reduction and cysteine alkylation may not be essential to achieve a good proteome coverage. This would therefore eliminate the exposure of the peptides to highly reactive alkylating reagents [30]. Furthermore, chemically labile amino acid residues may undergo sample preparation induced oxidation, deamidation, pyroglutamate formation, dehydration or excessive metal ion adduct formation [31–33]. Many of those modifications (or adducts) may, however, also occur as consequence of the natural protein ageing processes within the cell and are, therefore, difficult to discriminate from sample preparation artefacts [34].

A systematic evaluation – using a matrix-like approach, which leaves out one chemical at a time – and which investigates the impact on i) the % of identifications, ii) proteome coverage, iii) GO-term distribution and iv) post-isolation modifications, has not been performed for an unicellular organism to date. We therefore constructed a matrix of conditions, which include steps frequently employed in whole cell lysate proteomics, and applied it to the preparation of well-controlled chemostat grown yeast cells. The outcomes of the proteomics experiments were compared for their obtained proteome and amino acid sequence coverage and for their GO-term profiles. Moreover, we performed an unrestricted modification search using TagGraph, to identify reagent-induced modifications. Finally, we investigated the quality of the unidentified spectra using *de novo* peptide sequencing. In summary, this study provides a systematic evaluation of sample preparation protocols for the increasingly studied model organism and cell factory yeast. Thereby, we demonstrate the large impact of the different sample preparation procedures on proteome coverage and overall identification rates.

Ultimately, the performed study and publicly available large proteomics datasets, provide a valuable resource to select for the most suitable sample preparation elements for different experiments, and support the analysis of (native) peptide modifications.

## EXPERIMENTAL SECTION

### Yeast strain, growth media and storage

In this study, we used the minimal glycolysis (MG) yeast strain IMX372 (*MATa ura3-52 his3-1 leu2-3,112 MAL2-8c SUC2 glk1::SpHis5, hxk1::KlLEU2, tdh1::KlURA3, tdh2, gpm2, gpm3, eno1, pyk2, pdc5, pdc6, adh2, adh5, adh4*), which under the selected growth conditions shows now phenotypic alterations compared to the parent CEN.PK lineage [35, 36]. Shake flask and chemostat cultures were grown in synthetic medium (SM) containing 5.0 g·L^-1^ (NH4)_2_SO_4_, 3.0 g·L^-1^ KH_2_PO_4_, 0.5 g·L^-1^ MgSO_4_·7H_2_O and 1 mL·L^-1^ trace elements in demineralized water. The medium was heat sterilized (120°C) and supplemented with 1 mL·L^-1^ filter sterilized vitamin solution [37]. For shake flask cultures 20 g·L^-1^ heat sterilized (110°C) glucose (SMG) was added. In chemostat cultures, 7.5 g·L^-1^ glucose was added and the medium was supplemented with 0.2 g·L^-1^ antifoam Pluronic PE 6100 (BASF, Ludwigshafen, Germany). Frozen stocks of *S. cerevisiae* cultures were prepared by the addition of glycerol (30% v/v) in 1 mL aliquots for storing at -80°C.

### Yeast chemostat cultures and sampling

Aerobic shake flask cultures were grown at 30°C in an Innova incubator shaker (New Brunswick™ Scientific, Edison, NJ, USA) at 200 rpm using 500 mL round-bottom shake flasks containing 100 mL medium. Duplicate aerobic chemostat cultures were performed in 2 L laboratory fermenters (Applikon, Schiedam, The Netherlands) with a 1 L working volume in duplicate. SM-medium was used and maintained at pH 5 by the automatic addition of 2 M KOH. Mixing of the medium was performed with stirring at 800 rpm. Gas inflow was filter sterilized and compressed air (Linde Gas, Schiedam, The Netherlands) was sparged to the bottom of the bioreactor at a rate of 500 mL·min^-1^. Dissolved oxygen levels were measured with Clark electrodes (Mettler Toledo, Greinfensee, Switzerland). The temperature of the fermenters was maintained at 30°C. The reactors were inoculated with exponentially growing shake flask cultures of *S. cerevisiae* strain IMX372 to obtain an initial optical density (OD_660_) of approximately 0.4. Following the batch phase, the medium pump was switched on to obtain a constant dilution rate of 0.10 h^-1^. Chemostat cultures were assumed to be in steady state when, after five volume changes, the culture dry weight, oxygen uptake rate and CO_2_ production rate varied less than 5% over at least 2 volume changes. The dilution rate and carbon recovery were determined after each experiment.

### Analytical methods

OD_660_ measurements to monitor growth were performed on a JENWAY 7200 spectrophotometer (Cole-Parmer, Stone, UK). The biomass dry weight was determined in duplicate by extracting 10 mL of broth and filtrating it over a filter with 0.45 μm Ø pores while a vacuum was applied to the filter. The filters were washed twice with 10 mL of demineralized water. Prior to use, the filters were dried in the oven for at least 24 hours at 70°C. After filtration, the filters were dried in a microwave oven at 360 W for 20 min leaving only dry biomass. For extracellular metabolite determinations, broth samples were centrifuged for 5 min at 13.000 g and the supernatant was collected for subjection to a Waters alliance 2695 HPLC (Waters Chromatography B.V., Etten-Leur, The Netherlands) with an Animex HPX-87H ion exchange column (Bio-Rad, Hercules, CA, USA). The HPLC was operated at 60°C and 5 mM of H_2_SO_4_ was used as mobile phase at a rate of 0.6 mL·min^-1^. Off-gas concentrations of CO_2_ and O_2_ were measured using an NGA 2000 analyzer. Proteome samples (1 mL, at approx. 3.6 g·L^-1^ dry weight) were taken from steady state cultures. The samples were collected in multifold in trichloroacetic acid (TCA) (Merck Sigma, Cat. No. T0699) with a final concentration of 10% or in five volumes of ice-cold methanol (MeOH) (Thermo Fisher, Cat. No. 15654570). Samples were centrifuged at 4000 g for 5 min at 4°C. Cell pellets were frozen at -80°C [11].

### Proteomics sample preparation protocols (an extended version of all sample preparation protocols are provided in the SI document)

Yeast cell culture pellets were resuspended in lysis buffer composed of 100 mM triethylammonium bicarbonate (TEAB) (Merck Sigma, Cat. No. T7408) containing 0.1%, 1% sodium dodecyl sulphate (SDS) (Merck Sigma, Cat. No. L4522) or 8 M Urea (Merck Sigma, Cat. No. U5378) and phosphatase/protease inhibitors. Yeast cells were lysed by glass bead beating and thus shaken 10 times for 1 minute with a bead beater alternated with 1 minute rest on ice. For in-solution methods, proteins were reduced by addition of 5 mM DTT (Merck Sigma, Cat. No. 43815) or 5 mM TCEP (Merck Sigma, Cat. No. C4706) and incubated for 1 hour or 30 min at 37°C or 56°C, respectively. Subsequently, the proteins were alkylated for 30 min or 1 hour at room temperature in the dark by addition of 15 mM iodoacetamide (Merck Sigma, I1149) or 50 mM acrylamide (AA) (Merck Sigma, Cat. No. A9099), respectively. Protein precipitation was performed by addition of four volumes of ice-cold acetone (−20°C) (Merck Sigma, Cat. No. 650501) or TCA to a final concentration of 20% and proceeded for 1 hour at -20°C or 30 min at 4°C, respectively. The proteins were washed twice with acetone and subsequently solubilized using 100 mM ammonium bicarbonate (ABC) (Merck Sigma, Cat. No. 09830). Alternatively, for filter-aided sample preparation (FASP), proteins were loaded to a filter (Merck-Millipore, Microcon 10 kDa, Cat. No. MRCPRT010) after bead beating and reduced by addition of DTT and alkylated with iodoacetamide, as described earlier. After alkylation, proteins were washed four times with TEAB and ABC buffers. For all protocols, proteolytic digestion was performed by Trypsin (Promega, Cat. No. V5111), 1:100 enzyme to protein ratio (v/v), and incubated at 37°C overnight. For filter-aided sample preparation protocols, peptides were eluted from the filters after digestion using ABC and 5% acetonitrile (ACN) (Thermo Fisher, Cat. No. 10489553) / 0.1% formic acid (FA) (Thermo Fisher, Cat. No. 10596814) buffers consecutively. For all protocols, solid phase extraction was performed with an Oasis HLB 96-well μElution plate (Waters, Milford, USA, Cat. No. 186001828BA). Peptide fractions were eluted using MeOH buffer containing trifluoroacetic acid (TFA) (Merck Sigma, Cat. No. 302031), FA or ABC. Eluates were dried using a SpeedVac vacuum concentrator. Dried peptides were resuspended in 3% ACN / 0.01% TFA prior to MS-analysis to give an approximate concentration of 500 ng per μL.

### Shotgun proteomic analysis

For each protocol an aliquot corresponding to approx. 750 ng protein digest was analysed using a one-dimensional shotgun proteomics approach [38]. Each sample was analysed in two technical replicates. Briefly, the samples were analysed using a nano-liquid-chromatography system consisting of an EASY nano LC 1200, equipped with an Acclaim PepMap RSLC RP C18 separation column (50 μm x 150 mm, 2 μm, Cat. No. 164568), and a QE plus Orbitrap mass spectrometer (Thermo Fisher Scientific, Germany). The flow rate was maintained at 350 nL·min^-1^ over a linear gradient from 5% to 30% solvent B over 90 min, then from 30% to 60% over 25 min, followed by back equilibration to starting conditions. Data were acquired from 5 to 120 min. Solvent A was H_2_O containing 0.1% FA, and solvent B consisted of 80% ACN in H_2_O and 0.1% FA. The Orbitrap was operated in data-dependent acquisition (DDA) mode acquiring peptide signals from 385–1250 m/z at 70,000 resolution in full MS mode with a maximum ion injection time (IT) of 100 ms and an automatic gain control (AGC) target of 3E6. The top 10 precursors were selected for MS/MS analysis and subjected to fragmentation using higher-energy collisional dissociation (HCD). MS/MS scans were acquired at 17,500 resolution with AGC target of 2E5 and IT of 75 ms, 2.0 m/z isolation width and normalized collision energy (NCE) of 28.

### Mass spectrometric raw data processing. *De novo* sequence analysis

*De novo* sequencing was performed using the algorithm available via PEAKS Studio X+ (Bioinformatics Solutions Inc., Waterloo, Canada) [39], allowing 10 ppm parent ion and 0.5 Da fragment ion mass error, and oxidation as variable modification, where the resulting *de novo* sequences were exported to ‘de novo peptide.csv’ files for further unrestricted modification search using TagGraph, as described below.

### Taxonomic profiling

Taxonomic purity assessment using the same *de novo* peptide sequences was performed as described recently by H.B.C. Kleikamp et al. (2021) [40].

### Database searching

Database searching against the proteome database from *S. cerevisiae* (Uniprot, strain ATCC 204508 / S288C, Tax ID: 559292, June 2020, excluding 13 glycolytic isoenzymes) was performed using PEAKS Studio X+, allowing for 20 ppm parent ion and 0.02 m/z fragment ion mass error, 3 missed cleavages, carbamidomethyl or acrylamide as fixed (or none), and methionine oxidation and N/Q deamidation as variable modifications. To control false-positive peptide identifications, a uniform 1% false discovery rate (FDR) was applied to peptide spectrum matches (PSM), and subsequently the protein identifications required ≥2 unique peptides. Results from the PEAKS DB search were exported to ‘proteins.csv’ and ‘DB search psm.csv’ files, containing the identified proteins and identified DB search peptide-spectrum matches, respectively

### Unrestricted modification search

TagGraph [41] was used to perform unrestricted global peptide modification search using the (mzML-formatted) mass spectrometric raw data and the *de novo* sequences obtained from PEAKS Studio, using the yeast proteome database plus the CRAPome contaminant sequences [42]. The analysis was performed allowing for 10 ppm precursor mass tolerance, cysteine carbamidomethylation or acrylamidation as static modifications, and methionine oxidation as a differential modification, as described by Devabhaktuni et al. (2019) [41]. TagGraph.1.8 was installed on a Windows desktop Docker container, and the processing of multiple files was automated via a PowerShell script. An FDR of 1%, 10 ppm parent ion and a maximum absolute deviation of 0.1 Da between experimental and database modification mass were applied to the analysis results and these were exported to ‘.txt’ files for further analysis.

### Interconversion of mass spectrometric raw data

Conversion of the mass spectrometric raw data was performed using peak picking ‘vendor’ into ‘.mzML’ and ‘.mzXML’ using the msConvertGUI tool (ProteoWizard) [43].

### Identification of scans originating from glycopeptides

First, the fragmentation spectra were searched for the presence of glycan-typical HexNAc oxonium fragment ions (204.087, [M+H^+^]^+^) using functions from the Matlab ‘sugar miner’ script as described recently [44]. Those scans indicate glycopeptides–which remain unidentified by the chosen database, or open modification search parameters. Scans with strong oxonium ion signals were summarized in an ‘.xlsx’ table.

### Data processing and visualization. Overall number of modified peptides and types of modifications

The results of the unrestricted modification search using TagGraph were used to determine the overall volume as well as the types of modifications found after applying the different protocols. First, the mass shifts (=deviations from the unmodified peptide mass) were collected from the “.txt” files and binned using the Matlab ‘histcounts’ function, at a bin width of 0.01 Da. This procedure was done for the combined dataset as well as for each protocol separately. Mass shifts with more than 5 occurrences per averaged conditions were tested for significant changes across all protocols using Matlabs ‘anova1’ function. Mass shifts with significant changes (p<0.01) across all conditions (2 biological and 2 technical replicates) were visualized by Euclidean distance clustering using the Matlab ‘clustergram’ function, standardizing along the rows (mass shifts) of data, clustering along the columns of data, and then clustering along the rows of row-clustered data. Default color variation has been used which shows for values between -3 and 3, where values above and below show the same maximum color tone.

### Spectral Average Local Confidence (ALC) score histograms

After the scan numbers of each protocol were allocated to different categories, the Average Local Confidence (ALC) score distribution for identified and unidentified MS/MS scans was determined to evaluate the quality of the unidentified spectra. The ALC score was extracted from the *de novo* sequences ‘.csv’ files and were therefore only determined for the spectra that were *de novo* sequenced. The ALC scores of the identified and unidentified scans were binned separately using the ‘histcounts’ function in Matlab. The resulting distributions were plotted as bar graph and exported as table.

### Proteome and amino acid sequence Coverage

The proteome coverage was determined using the PEAKS-DB search results ‘proteins.csv’. For this, the number of proteins per protocol was calculated from proteins with >2 unique peptides per protein. The percentage of observed proteins for each protocol was subsequently calculated by normalizing to the number of total proteins in the *S. cerevisiae* (Uniprot) protein sequence database. The average protein sequence coverage per protocol was further extracted from the PEAKS ‘proteins.csv’ output files. The sequence coverage (%) was moreover used to determine the amino acid coverage, in which the sequence coverage of each identified protein was summed and related to the total number of protein sequences present in the sequence database. The average for the proteome and amino acid coverage was calculated for each biological replicate and the deviations from the average for each protocol were plotted as bar graphs. A two-tailed unpaired Student’s t-test was performed to determine if the relative coverage change between protocols was statistically significant.

### Ontology analysis

The ‘proteins.csv’ files obtained through PEAKS-DB were used to determine the differences in the cellular component distribution. Python 3.8 [45] was used to programmatically link the Uniprot accession numbers of the identified proteins (with 2 unique peptides) to the Gene Ontology (GO) [46] terms using Retrieve/ID Mapping function on Uniprot [47]. The GO library was imported using the ‘goatools’ module in Python to retrieve the cellular component terms in the GO hierarchy. Functions of the proteins were summarized in pie charts and bar graphs based on their cellular component GO terms. GO terms with less than one percentage contribution were clustered into category “other”.

### Overall % of identified fragmentation spectra

The overall number of identified fragmentation (MS/MS) spectra for each protocol was determined by combining the outputs obtained from TagGraph, PEAKS database search and the sugar oxonium ion search. First, scan numbers that did not result into amino acid sequence candidates by PEAKS *de novo* sequencing were allocated to the ‘No ALC’ category. Next, identifications were extracted from the TagGraph files. Thereby, identified scans were allocated to two categories, ‘unmodified peptides’ and ‘modified peptides ‘. The scan numbers of the peptides identified with the second search engine were extracted from the PEAKS-DB search peptide-spectrum matches files ‘DB search psm.csv’. The scan numbers were compared with the scans of the *de novo* sequences and TagGraph results. Identifications that were only found with PEAKS database search, were allocated to the ‘second search engine’ category. The category “sugars” represents the scans that were identified as potential glycopeptides. If scans were present in multiple categories, including TagGraph, the scans were allocated to TagGraph identifications (modified or unmodified peptides). If scans were identified containing sugar fragments and by PEAKS database search, the scans were allocated to PEAKS-DB search identifications. Furthermore, ‘No ALC’-allocated scans that could be identified containing sugar fragments or via PEAKS database search, were allocated to one of the respective categories. Finally, the MS/MS scans that could not be identified by any of the before mentioned categories, were allocated to the ‘unidentified scans’ category. As final check, the summed scans of all established categories required to be equal to the sum of MS/MS scans in the raw mzXML files. The distribution of the scan identifications amongst different categories was visualized using stacked bar graphs, in which the number of scans in each category was normalized against the total number of MS/MS scans in percentages, for each protocol.

### Data availability

Mass spectrometric raw data, protein sequence database, and search files have been deposited at ProteomeXchange server and are publicly available under the project code PXD026806.

## RESULTS AND DISCUSSION

### Comparison of yeast whole cell lysate sample preparation protocols

A systematic comparison of a matrix of different sample preparation procedures was performed to investigate the impact on spectral identification rates and quality, achieved proteome coverage, amino acid sequence coverage, GO-terms distribution and reagent-induced peptide modifications (Table 1, Figure 1). Duplicate aerobic chemostat cultures of the IMX372 *S. cerevisiae* strain [36] were cultured in glucose-limiting conditions (Supplementary Table S1), which provided highly reproducible yeast cell biomass. Proteome samples were taken during steady-state conditions from both biological replicates, in which the cellular metabolism was quenched using ice-cold TCA or methanol to preserve the proteome and associated post-translational modifications. Cell lysis was performed with SDS- or Urea-containing buffers employing bead-beating in all cases. Crude cell lysates were then treated with an in-solution or FASP approach, in which proteins were reduced with TCEP or DTT and alkylated with iodoacetamide, acrylamide or alternately, were left untreated. For in-solution digestion methods, proteins were precipitated with acetone or TCA. After overnight digestion with trypsin, solid-phase extraction was performed to remove contaminants by SPE using Oasis HLB cartridges, which make use of a co-polymer of divinylbenzene and vinyl pyrrolidinone that shows an enhanced retention of polar peptides (Waters). Elution from the cartridges was achieved with a variety of buffers to assess the influence of different additives (Table 1). Finally, the samples of the different protocols were analysed using a (short) one-dimensional gradient with duplicate injections.

**Table 1:**
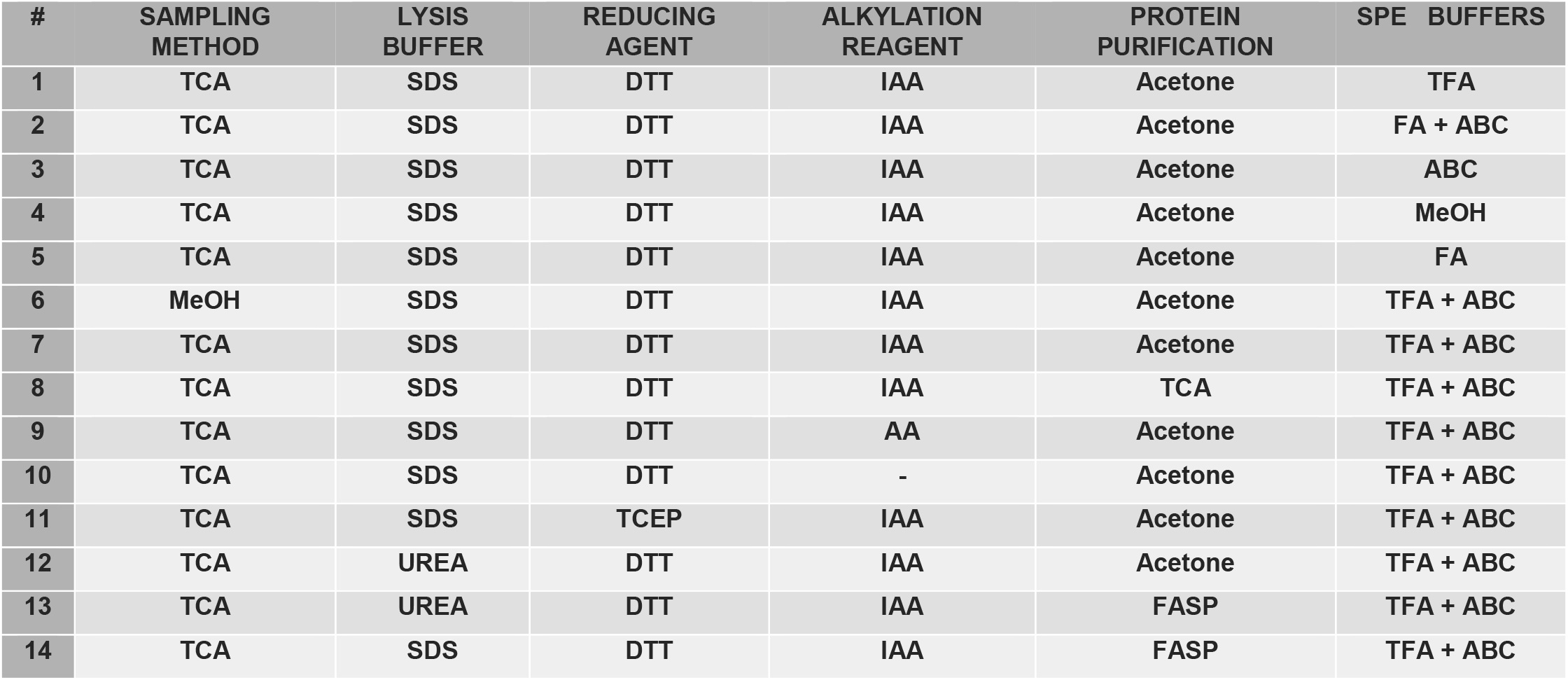
Matrix of investigated sample preparation protocols (1–14). Abbreviations: SPE, solid phase extraction; TCA, trichloroacetic acid; SDS, sodium dodecyl sulphate; DTT, Dithiothreitol; TCEP, Tris(2-carboxyethyl)phosphine; IAA, iodoacetamide; AA, acrylamide; TFA, trifluoroacetic acid; FA, formic acid; ABC, ammonium bicarbonate. The detailed protocols are provided in the S.I. materials.

**Figure 1:**
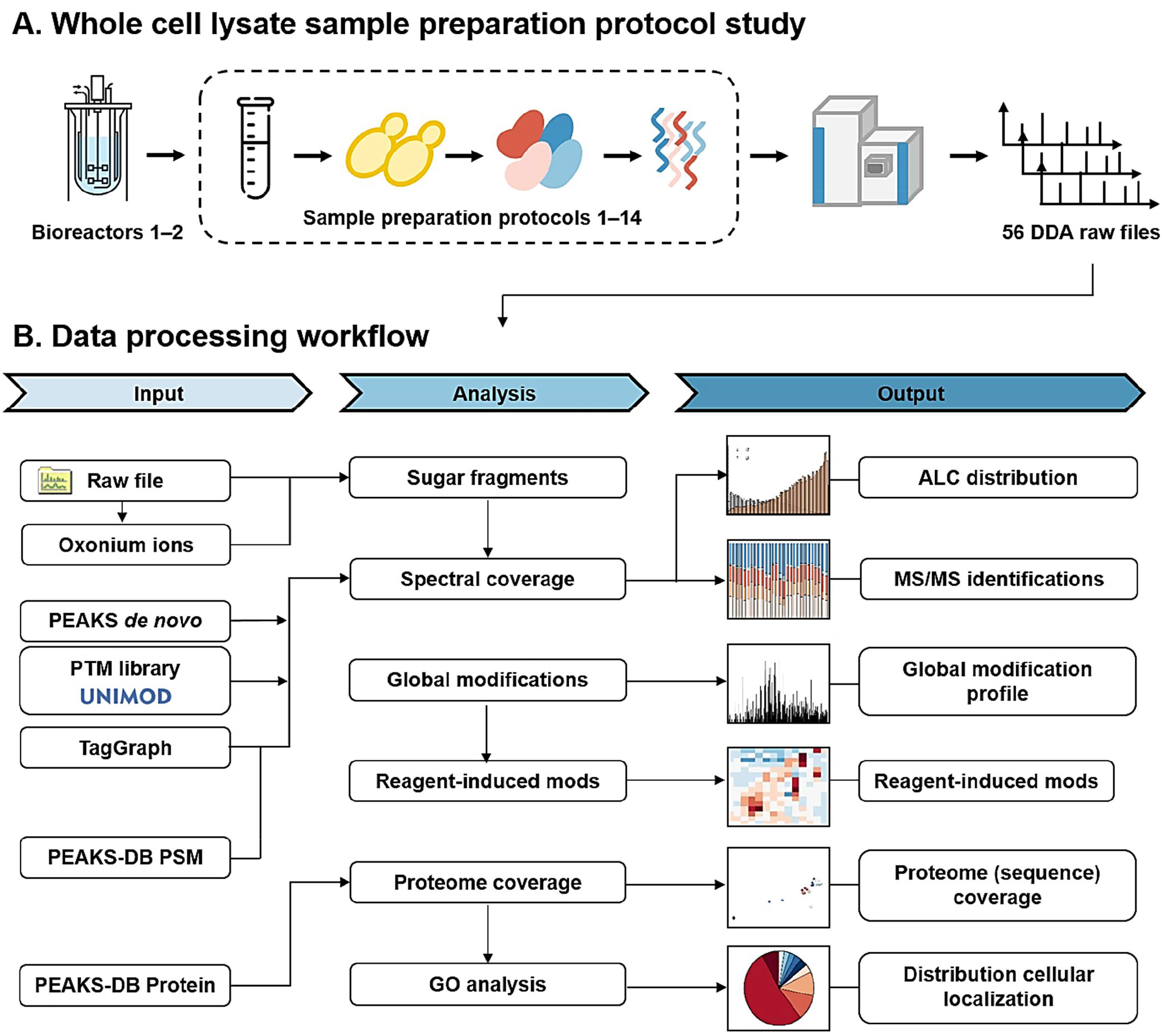
Whole cell lysate sample preparation study for yeast and established data processing pipeline. A. The employed study performed yeast cultivation in aerobic chemostats, sample collection and quenching, cell lysis by bead beating, protein extraction from cells and subsequent reduction and alkylation of the proteins. Proteins are then concentrated using protein precipitation or a filter-aided approach. Enriched proteins are subsequently digested using Trypsin, where the obtained peptides are subjected to solid phase extraction (SPE) to remove contaminants prior to shotgun proteomic analysis. **B**. The obtained MS/MS spectra were *de novo* sequenced using PEAKS and subsequently identified through database (PEAKS-DB) and unrestricted modification searches (TagGraph). A data analysis pipeline was established to determine the % of identified spectra, spectral quality, proteome and amino acid sequence coverage. Peptides identifications obtained by TagGraph were furthermore used to determine the modification profiles for every protocol. Finally, proteome were annotated with GO-terms to investigate the distribution according to ‘cellular components’.

On average, 35420±5909 (biological replicate 1) and 37261±8271 (biological replicate 2) MS/MS scans were obtained across all protocols (Figure 2, Supplemental Figure S1). Furthermore, the analysis of the protocols resulted in 23631±5237 and 23477±6590 peptide spectrum matches (PSMs) (Figure 2a), resulting in 66% and 61% MS/MS spectrum identifications on average (Figure 2b). However, protocols 8 and 14 resulted consistently in a considerable lower number of MS/MS spectra for both biological replicates, albeit that a comparable amount of proteolytic digest (750 ng) was analysed. For protocol 8, protein precipitation was performed with TCA/acetone instead of acetone solely. Difficulties in re-dissolving the protein pellet can arise for the TCA/acetone precipitation approach and may ultimately impact the overall recovery. It therefore becomes necessary to use stronger buffers, larger volumes and additional mechanical disruption of the pellet, to solubilise the protein [15, 18, 48]. In this study, protein pellets were dissolved in ammonium bicarbonate buffers to allow subsequent trypsin digestion, which did not prove difficult for acetone-precipitated samples. However, the TCA/acetone procedure led to a partially insoluble pellet, and a considerably lower number of proteins were in the following identified, compared to when only using acetone-precipitation (20% vs. 29% proteome coverage for protocol 8 = TCA/acetone, vs protocol 7 = acetone; Figure 2c). Similar results were observed in other studies, which was attributed to the increased protein denaturation caused by this approach [15, 49] [50] [51]. Samples treated according to protocol 14 were subjected to FASP using an SDS containing buffer. SDS is known to interfere with binding and elution during the reverse-phase separation of the peptides and severely suppresses ionisation by electrospray ionisation [52–54]. Multiple rounds of centrifugation were applied to ensure SDS removal, however, residual SDS may have still impacted the LC-MS/MS analysis, resulting in a lower total number of MS/MS scans (13232 MS/MS spectra on average for protocol 14, vs. 36340 MS/MS spectra on average across all protocols). However, an earlier study buffer exchanged SDS with Urea following solubilisation of the proteins, allowing for high protein identification rates, because SDS was likely successfully withdrawn from the sample [19]. Hence, FASP sample preparation in combination with SDS-containing buffers is not recommended unless the complete removal of SDS can be ensured.

**Figure 2:**
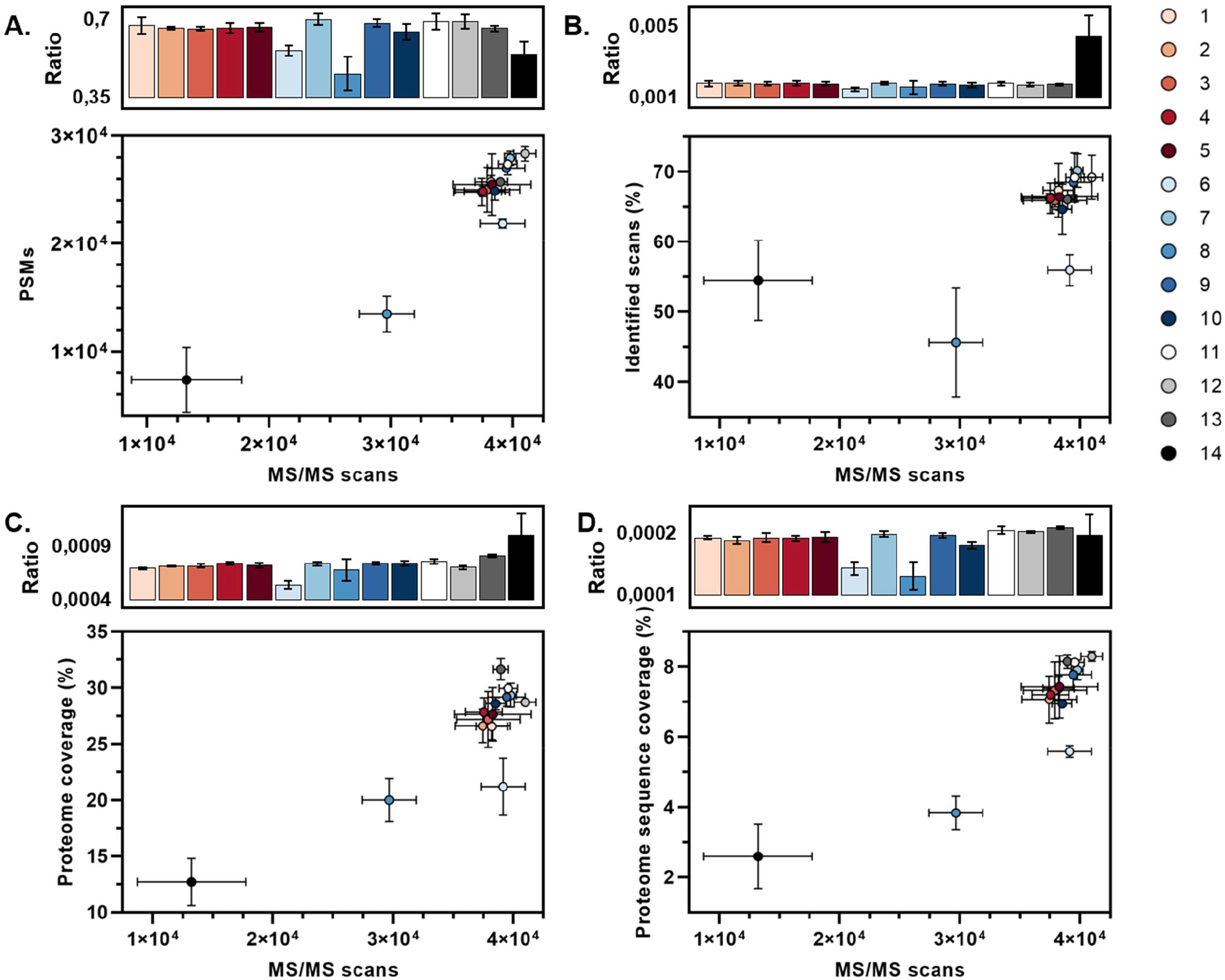
Achieved % of identified fragmentation spectra, proteome and sequence coverage for the different sample preparation protocols. The number of Peptide Spectrum Matches (PSMs) (**A**); identified scans (%) (**B**); proteome coverage (%) (**C**) and proteome sequence coverage (**D**), were plotted against the number of MS/MS scans obtained per protocol. Each coloured circle represents the average of averages of two biological replicates with each 2 technical replicates (2×2), while the standard deviation is represented by the error bars. In addition, the bars depicted above each plot show the ratios of the acquired PSMs, identified scans and proteome (sequence) coverage vs. MS/MS scans for each protocol. The ratios are averages of two biological replicates with each 2 technical replicates (2×2), while the standard deviation is shown as error bars. The proteome coverage was calculated based on the identified proteins per protocol as a percentage of the total number of proteins in yeast (known ORFs). In addition, the proteome sequence coverage was calculated based on the identified amino acids in the proteomics experiments, as a percentage of the total proteome amino acid sequence (sequence coverage).

### Overall proteome coverage is strongly affected by sample preparation procedures

MS/MS spectra were identified using database searching employing common statistical filtering criteria (see materials and methods) and requiring at least 2 unique peptides per protein identification. The proteome coverage was calculated based on the proteins identified per protocol as a percentage of the total number of proteins in yeast (known ORFs). An average proteome coverage of approximately 26% and 27% was achieved amongst the different methods for biological replicates 1 and 2, respectively (Figure 2c). Small differences in the average depth of the proteome coverage between both biological replicates were observed, most likely because the experiments were conducted at different time periods and, therefore, the instrumental performance may have been slightly different. Three protocols (6, 8 and 14) differed substantially from the other procedures. As discussed above, for protocols 8 and 14, LC-MS/MS analysis was likely compromised by incompatibility with the used reagents. Protocol 6, on the other hand, resulted in an average number of 39152 MS/MS spectra, which is slightly higher than the average of 36341 MS/MS spectra across all protocols (Figure 2b). Since the total ion count was very comparable, its suggestive that a similar amount of proteolytic digest was injected to the LC-MS system. This was the only sample that was subjected to quenching with methanol as opposed to TCA during sampling. Methanol quenching (applied to yeast) might, therefore, significantly impact the number of identifiable proteins. TCA as a quenching solution has been also proven useful in quantitative phosphoproteomics measurements recently [11]. Methanol, on the other hand, is routinely employed to rapidly arrest the cellular metabolism when performing metabolomics studies [55]. Methanol has been also used to co-extract metabolites and proteins from yeast, only recently [56]. In the present study, the methanol quenching bias was observed consistently for all biological and technical replicates. Protein aggregation due to exposure to methanol followed by poor resolubilisation is likely the cause for the low proteome coverage, however, the exact mechanism remains to be explored.

Nonetheless, the highest proteome coverage (31% and 32%, for biological replicates 1 and 2, respectively) was obtained with sample 13, in which a filter-aided approach was used in combination with Urea buffer. Similar results have been found in previous studies, in which FASP outperformed in-solution approaches in terms of the number of protein identifications [57, 58].

The proteome coverage does not seem to have been significantly impacted by the type of detergent used during lysis (SDS or Urea) for samples 7 and 12, respectively. Typically, SDS is used as an ionic and denaturing surfactant that can disrupt cell membranes and cause protein denaturation by disrupting protein–protein interactions. Urea, on the other hand, is a chaotrope that can bind to proteins, thereby causing protein unfolding [59]. Following protein extraction, protein reduction was performed by either TCEP or DTT. The type of reducing agent used during sample processing was also not a key determinant in the outcome of the proteome analysis. This was very similarly observed for yeast in another study [13].

In this study, proteins were subsequently alkylated with iodoacetamide, acrylamide or alkylation was left out completely (samples 7, 9 and 10, respectively). Expectedly, and in line with a recent publication [30], the absence of an alkylation step dramatically reduced the detection of cysteine-containing peptides, although the depth of the proteome coverage remained nearly unchanged. However, the number of MS/MS scans and obtained peptide spectrum matches, as well as sequence coverage, was steadily lower for the non-alkylated sample (protocol 10), when compared to the alkylated samples (protocols 7 and 9). Furthermore, no significant changes in the number of identified proteins could be observed for the different solvents used for solid-phase extraction (protocols 1– 5, 7, Figure 2 and Supplemental Table S2 and S3). Though, the number of MS/MS scans, identified proteins and proteome (sequence) coverage was consistently higher for protocol 7 (Figure 2, Supplemental Table S2 and S3), in which a combination of basic and acidic MeOH buffers were used for elution, thereby maximising the recovery of peptides across a large pH range. Moreover, TFA (protocol 7) appeared to be a better choice for peptide elution compared to formic acid (FA) (protocol 2 and 5) because the proteome (sequence) coverage was repeatedly higher when using TFA compared to FA (Supplemental Table S2 and S3).

A gene ontology (GO) analysis [46] was performed to investigate whether the different protocols bias towards specific cellular components, such as cytosolic soluble proteins, or oppositely, towards hydrophobic membrane proteins. Therefore, GO terms were assigned to proteins based on their ‘cellular component’. Thereby, the distribution of the cellular localisation of the observed proteins was comparable across the different protocols, which is exemplified in Figure 3a for protocol 13 (biological replicate 2, Supplemental Table S4). Approximately 53% accounted for the intracellular fraction (GO-term ‘organelle’), whereas ∼11% of the identifications belonged to membrane proteins. Yet, a slight increase in the average relative value of membrane proteins was observed for protocols 8 and 14 (Figure 3b). Irrespective of the protocol, proteins were extracted using bead beating, which is a relatively harsh and likely reproducible approach. Nevertheless, a recent study demonstrated that extraction methods can significantly contribute to the variability of proteomics experiments [8].

**Figure 3:**
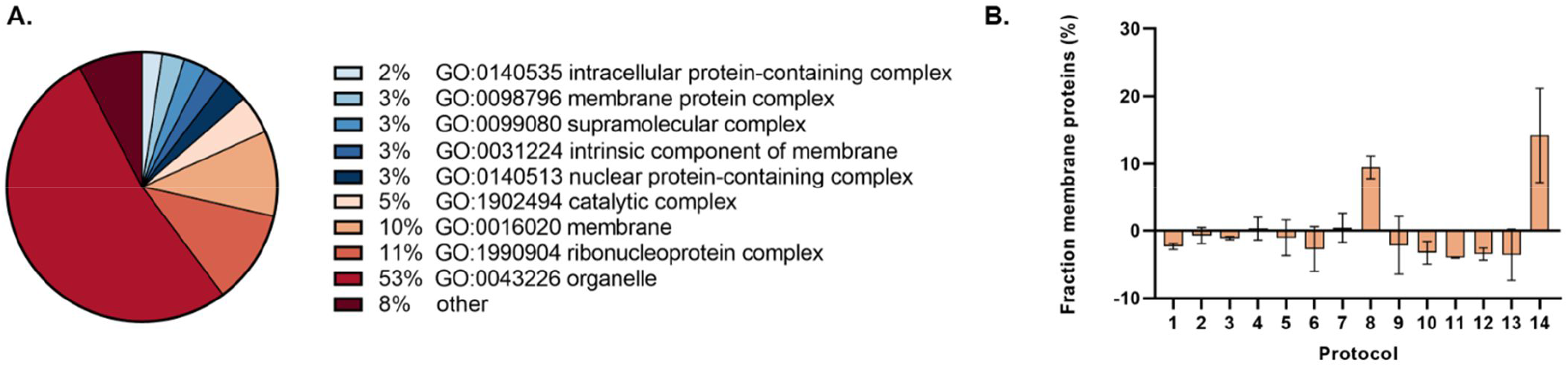
Distribution of Gene Ontology (GO) terms for identified proteins. **A)** Based on the classifications of GO annotation, the identified proteins (with at least 2 unique peptides) were categorized into cellular components, which is displayed in the pie chart for protocol 13 (biological replicate 1). Categories with GO terms less than 1% are classified as ‘other’. Here, the average of two biological replicates with each 2 technical replicates (2×2) is shown. **B)** The average fraction of identified membrane proteins across all protocols was 11%. The deviations from the average are shown in percentage for each protocol (1–14).

Commonly, proteins are already reported with a few peptides, e.g. 2 unique peptides. While acceptable for qualitative and quantitative analysis, further characterisation of individual proteoforms as well as analysis of post-translational modifications demand a higher sequence coverage. Therefore, we quantified the fraction of the total proteome amino acid sequence space covered for every protocol. An average proteome sequence coverage of 6.7% and 7.0% was observed across all protocols for biological replicates 1 and 2, respectively (Figure 2d). Hence, for the employed (relatively short) one-dimensional gradient—despite the good proteome coverage—a rather small fraction of the amino acid sequence space was actually observed in our experiments. In the present study, an average protein sequence of 26% and 25% was obtained amongst the identified proteins (Supplemental Figure S2). This value is proportional to the average amino acid sequence coverage of 28% obtained by Herbert et al. (2014), who obtained through extensive experiments a near-complete proteome coverage [61]. Nevertheless, the proteome coverage strongly depends on the protein amino acid sequences, and therefore, on the cleavage specificity of the employed protease(s) [60].

One of the challenges to improve proteome sequence coverage arises from signal suppression during electrospray ionisation (particularly for short one-dimensional gradients), where peptides with higher basicity tend to ionise preferably, and lower abundant peptides or highly hydrophobic or acidic peptides may remain undetected. Longer LC separations, additional peptide pre-fractionation, as well as 2-dimensional gradients significantly increases the number of identifications [62] [63–65]. Nevertheless, material requirements, sample processing and analysis time will increase proportionally, when using multi-dimensional chromatography or off-line peptide fractionation. Finally, multi-protease digestion has been demonstrated to boost not only the protein but also the proteome sequence coverage [66, 67].

As observed for the proteome coverage, protocols 6, 8 and 14 resulted in a much lower proteome sequence coverage (Figure 2d, Supplemental Table S2 and S3). Overall, while lysis buffer, reducing agent or SPE elution buffers seemingly did not impact proteome coverage, the alkylation step was critical for obtaining a high amino acid sequence coverage.

### The observed variability in peptide modifications

By using an unrestricted modification search (TagGraph) 221751 and 181496 mass shifts were observed across all used protocols, which derived from 991759 and 1043314 MS/MS spectra for biological replicates 1 and 2, respectively (Figure 4). A highly comparable mass shift profile was observed for both technical and biological replicates, confirming the reproducibility of the chemostat cultivation [68], the sample preparation protocols and the shotgun analysis. To discriminate between peptide modifications from biological (co/post-translational) and sample processing origin, we searched for alterations (or mass shifts) that were predominantly (or exclusively) present when using certain sample preparation protocols. Hence, we searched for mass shifts that showed a significant change in frequency (p<0.01) across all conditions and replicates, of which the averaged occurrences across all protocols is shown in Figure 5 (Supplementary Table S5). Albeit we employed combinations of well-established sample preparation steps, we observed a distinct number of condition-specific modifications, of which some had not been linked to sample preparation artefacts before.

**Figure 4:**
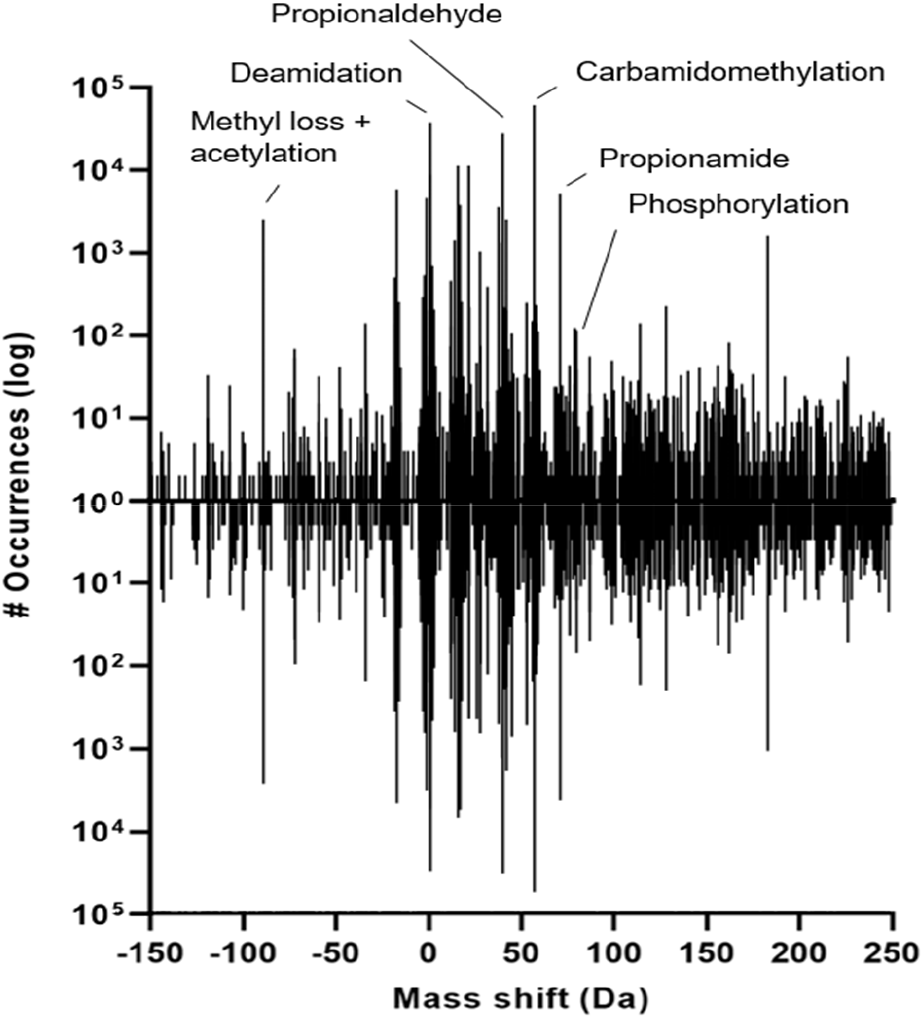
Sum of observed mass shifts in the different protocols using an unrestricted modification search approach (TagGraph). The number of occurrences corresponds to the number of peptide spectrum matches containing the mass deviation (log scale) for biological replicate 1 (upper histogram) and 2 (lower histogram). The total of both biological replicates show a highly comparable mass-shift profile.

**Figure 5:**
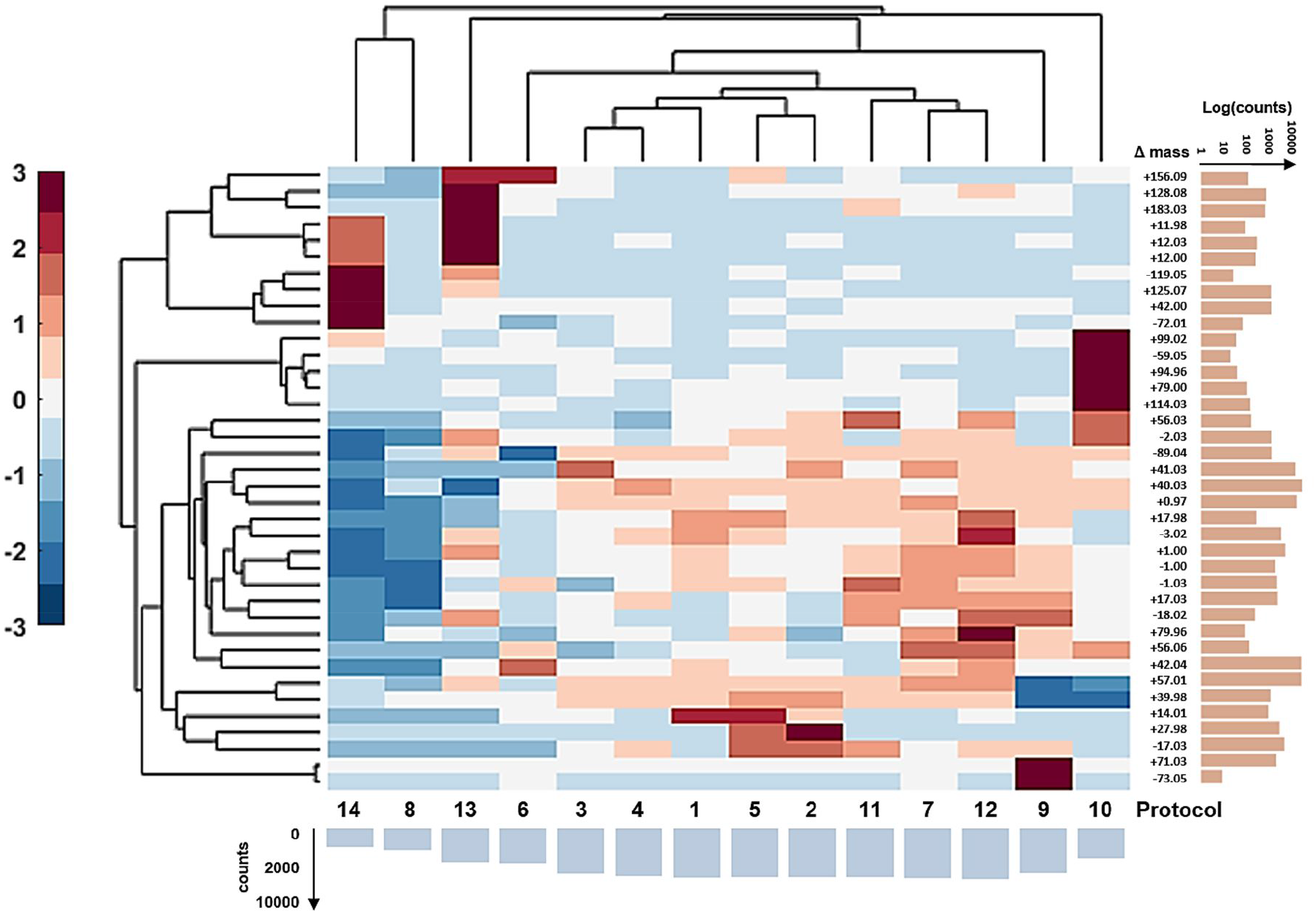
Clustered heat map of most prominent modifications identified by TagGraph across the different sample preparation protocols. Dendrogram and standardized clustered heat map of the relative change (%) of mass shifts observed across the sample preparation protocols. The mass shifts found for each protocol originating from each technical and biological replicates (2×2), were averaged per sample and normalized to the number of obtained MS/MS spectra. The bars below the heat map show the total counts of modifications observed for each sample and the bars on the y-axis show the total counts of the respective mass shift (modification) across the investigated samples.

For example, two different approaches to quench the cellular metabolism were employed, using either TCA or methanol. A mass shift of -89.04 Da was prominent in TCA-arrested yeast cells, with the notable exception of protocol 14 (SDS/FASP protocol). Furthermore, we used reagents for cell lysis and protein denaturation such as SDS and Urea. Interestingly, we did not observe any mass shifts related to the use of those reagents. In aqueous solutions, Urea dissociates upon heating over time. One of its degradation products is iso-cyanate, which can react with the N-termini of proteins/peptides and at the side-chain amino groups of lysine and arginine residues, to mimic *in vivo* carbamylation (+43 Da). This reaction is enhanced by high temperatures; however, Urea solutions in our study were only heated to 37°C for 1 hour, whereas 56°C is commonly used for protein reduction with DTT or TCEP [69]. Furthermore, the use of ammonium-containing buffers (and the removal of Urea before over-night digestion) seemed to minimise protein carbamylation when using Urea solutions [70], which was also observed in our study. Therefore, the relatively mild procedure used by our protocols seemingly avoided carbamylation reactions taking place.

No significant mass shifts were observed between samples reduced with TCEP or DTT. This is in line with an earlier study demonstrating that both reducing agents resulted in the same number of protein identifications [12, 13]. Nevertheless, acrylamide appeared to be more efficient in combination with DTT, whereas iodoacetamide performed better with TCEP [12]. As expected, alkylation reagent-related mass shifts were the most abundant artefacts (e.g., +57.01 for IAA with 61582 and 53462 occurrences). However the frequently reported methionine alkylation and the subsequent neutral loss of -48 Da from the molecular peptide ion [M-48+H^+^]^+^, which is the loss of the side-chain from oxidised methionine (Mueller & Winter [12]), was rarely detected. Moreover, double alkylation with the characteristic mass delta of +114 Da was compared to counts for single alkylation (+57 Da) for all samples (on average) below <0.20%. That mass addition was also observed in the non-alkylated sample, suggesting that this mass shift resulted rather from ubiquitinylation (‘GlyGly’) and not from excessive alkylation. The modification analysis further indicated that the applied alkylation procedure was sufficiently mild (in particular when removing excess reagent before over night digestion) with only little off-target, and hardly any excessive, alkylation taking place. On the other hand, all protocols that used IAA showed additional +39.98 Da mass additions, which presumably derive from an N-terminal S-carbamoylmethyl-cysteine cyclisation product [28, 71, 72]. Acrylamide showed only some additional -73.05 Da mass shifts at low frequency, the mechanism of which, however remains to be investigated. Interestingly, the protocol (10) employing reduction but no alkylation resulted in an additional series of (albeit low frequent) mass shifts (such as -59.05, 79.9, 94.9, 99.0), which likely originate from reactions of the free sulfhydryl group with compounds (naturally) being present in cellular extracts. An unexpected observation was the very abundant mass addition of +40.03 Da (labelled in Unimod http://www.unimod.org/ as ‘propionaldehyde’). This mass shift was present in all protocols that employed protein precipitation with acetone, but was strongly reduced for protocols that used TCA with acetone wash, and which was absent in protocols using FASP. This mass shift was in a recent study related to acetone artefacts, which in fact can impact some 5% of the proteome (peptides with glycine as the second N-terminal amino acid) [17]. The 40 Da mass shift, however, can also be misinterpreted for an amino acid substitution from glycine to proline. Acetone modification involves aldimine formation between the ketone and the amino groups of a peptide, which is regarded as being acid labile and, therefore, rather rearranges to a more stable form (imidazolidinone), resulting in the observed +40.03 Da mass addition [17]. Interestingly, we observed some few mass additions of +41.03 Da, which may indicate the presence of the proposed aldimine intermediate. Other artefacts that presumably also derive from acetone, such as +98 Da and to a lesser extent +84 and +112 Da [73], were only observed at trace levels or were not present at all. However, it is suggested that those artefacts seemed to be more pronounced at elevated pH and, therefore, our protocols might have prevented their formation. Even though acetone may increase peptide complexity substantially, this procedure is still commonly used in sample preparations, likely because it efficiently removes organic compounds such as inhibitors or compromising detergents, such as SDS [48]. The +28 Da mass adduct was nearly exclusively found for protocols in which formic acid was used during solid-phase extraction, confirming that yeast (at the given growth conditions), showed hardly any double methylation events. Previous studies provided similar findings, thereby highlighting that the proteome analysis outcome can be strongly affected by the use of this reagent [25, 26]. A recent work demonstrated that these unwanted modifications can be largely avoided by processing the samples at low temperatures [25]. Another interesting observation in regard to solid-phase extraction was that the number of ammonium adductions (+17 Da) was hardly impacted by the nature of the buffers used to elute the peptides from the solid-phase extraction cartridges. Those adducts were supposedly carried over from the ammonium bicarbonate buffers, used for proteolytic digestion.

For the samples prepared with the alternative filter-aided protocols, a number of mass shifts were observed that were detected at a much lesser frequency in samples prepared by in-solution digestion (e.g., one protocol: +41.03, 42.00 or both protocols: -89.05, 11,0 and -2,0). On the other hand (albeit at low frequency), a series of mass modifications such as -119.0, -72.0, 11.98, 12.0, 42.0, 125.07, 128.07, 156.09 and 183.03 were increased or exclusively observed, when using filter-aided protocols.

### Considerable variations in the fraction of overall identified spectra observed for the different sample preparation protocols

The number of identified MS/MS spectra for each sample was determined by combining identifications from TagGraph, PEAKS database search and the sugar oxonium ion fragment search. First, the spectra were *de novo* sequenced, where a fraction of all spectra (2.6 % and 3.8%, on average, for replicates 1 and 2, respectively) did not provide any *de novo* spectra, which were further allocated to the ‘unsequenced’ category (‘no ALC score’, Figure 6, Supplemental Table S6). The *de novo* sequences were then used to perform an unrestricted peptide (modification) search using TagGraph, which resulted in the identification of 28% and 26% of unmodified and 17% and 14% of modified peptides, on average, for replicates 1 and 2, respectively (at a 1% peptide FDR). Moreover, PEAKS was used as an additional/orthogonal search engine to maximise identifications of unmodified peptides. Thereby, the number of confidently identified peptides considerably increased to 20% and 22%, on average, for replicates 1 and 2, respectively. Because different search engines employ individual approaches and scoring matrices, a combination of multiple search engines typically achieves significantly more matches [5, 7, 74–79]. The PEAKS database (PEAKS-DB) is a hybrid approach that combines elements of *de novo* sequencing (short sequence tag extraction) and database searching [80]. TagGraph, contrarily, is an unrestricted *de novo* sequence-tag approach employing fast string-based searches followed by a probabilistic validation model optimised for the identification of modified peptides [41]. Furthermore, spectra were screened for known carbohydrate fragments, which indicate a glycopeptide spectrum that would likely remain unidentified by any of the above-employed approaches and, therefore, were allocated to the category ‘sugars’ (0.8% and 0.6%, on average, for replicates 1 and 2, respectively). Finally, the remainder of the *de novo* sequences that could not be allocated to one of the aforementioned categories were categorised as ‘unidentified’. Here, approximately 31 and 34% remained unidentified for replicates 1 and 2, respectively.

**Figure 6:**
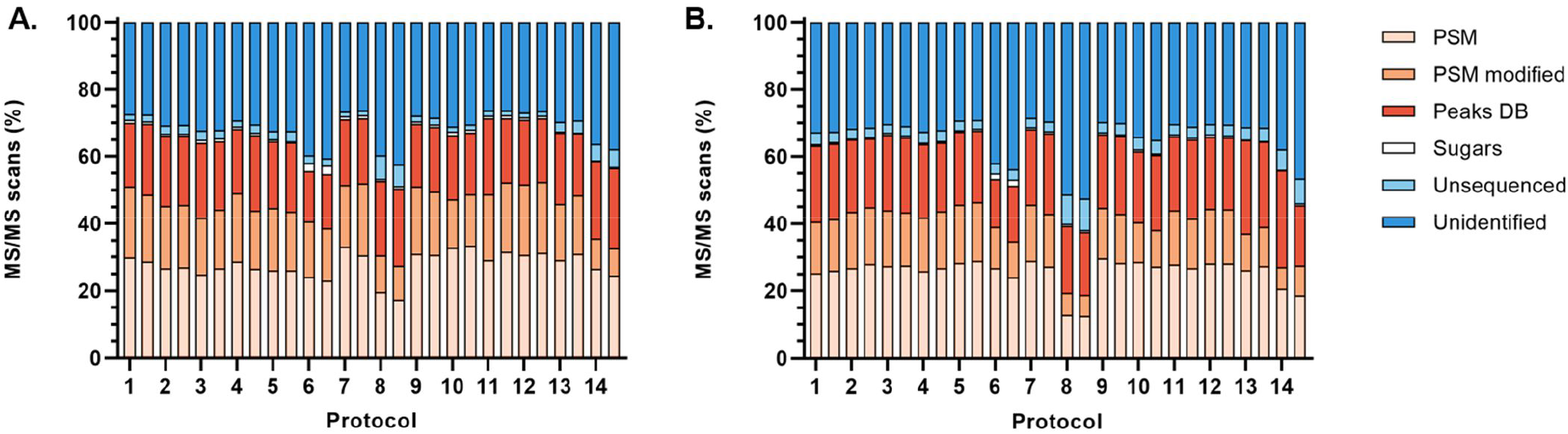
Overall spectral coverage obtained for the different sample preparation protocols. Allocation of total number of MS/MS spectra of each sample (1–14) into different categories: PSMs (TagGraph), PSMs modified (TagGraph), identified spectra by second search engine (PEAKS-DB), spectra containing oxonium fragments, ‘unsequenced’ and unidentified spectra for biological replicate 1 (**A**) and 2 (**B**).

As mentioned above, small differences in the identification rates between the two biological replicates were observed, presumably due to operational performance differences of the mass spectrometer at the different time periods. Still, a very similar pattern in allocation of the sequencing spectra is observed between the replicates (Figure 6). Expectedly, protocols 6 (methanol quenching), 8 (TCA precipitation) and 14 (FASP with SDS buffer) resulted in the lowest number of spectral identifications due to possible protein aggregation and following losses of protein during precipitation and LC-MS/MS incompatibility of reagents, respectively. Furthermore, the number of additional identifications with PEAKS-DB was considerably lower than for the other protocols when the samples were quenched with methanol. Conversely, a higher number of spectra from potential glycopeptides was observed using this protocol. This observation might be explained by the slightly increased fraction of membrane proteins (Figure 3), when using this protocol (number 6). The other protocols resulted in highly similar profiles and identification rates, with few exceptions. A considerably lower number of modified peptides was found for protocols 10 (no alkylation) and 13 (FASP, Urea). As speculated in the previous paragraph, this presumably results from the loss of cysteine-containing peptides, and acetone induced modifications for protocols 10 and 13. Finally, sample 13 (FASP with Urea buffer) resulted in one of the highest identification rates with a relatively low number of modified peptides. Nevertheless, a large fraction of all spectra remained unidentified.

### Investigating ‘the nature of the unidentified fraction’

As also observed in other studies [1, 2], approximately one third of the MS/MS spectra remained unidentified in the current study across all protocols. Recurrently, protocols 6, 8 and 14 led to a significantly higher fraction of unidentified spectra, on average, 44%, 54% and 45%, respectively. Other protocols obtained similar results, ranging from 30– 35% unidentified scans. Protocol 7 (in-solution, IAA) resulted in the highest number of identified spectra (70%), showcasing the key role of the type of protocol used on the identification rate. The inability to identify spectra can have various causes, such as impurities (or contaminations) in the sample, excessive co-fragmentation, poor quality spectra, unexpected modifications or peptide sequences not covered by the database (e.g., sequence variants and microbial impurities). First, the taxonomic purity of the samples was confirmed by a recently published *de novo* metaproteomic profiling approach [40], in order to exclude the presence of sample carry-over from foregoing analyses and to exclude the presence of bacterial contaminants. This was performed with protocol 13 for both biological replicates, for which no impurities could be identified (data not shown), confirming that no carry-over took place and that the yeast cultures were of highest purity.

A high quality of the fragmentation spectra is a prerequisite for confident peptide spectrum matching. Therefore, we assessed the quality by *de novo* sequencing and the associated average local confidence score (ALC), as determined by PEAKS [39]. In general, peptides with higher ALC scores indicate a more complete peptide fragment ion coverage and, therefore, are expected to have a higher chance of correct identification. As example, the ALC scores of the (un)identified spectra of biological replicate 1 processed with protocol 13 is visualised in Figure 7 (Supplemental Table S6). The ALC distribution is, as expected, very different for the fraction of identified and unidentified spectra. The ALC scores of the unidentified spectra spread predominantly around low numbers (where all identified spectra show very high ALC scores), suggesting that reduced quality (peptide fragment ion coverage) is the main reason for the lack of identification. Nevertheless, a small fraction of the unidentified spectra showed high ALC scores. Most of the *de novo* sequences were close to sequences in the yeast protein sequence database and may therefore represent sequence variants (e.g., single nucleotide polymorphisms SNPs, or isoform- and allele-specific variants of proteins) or additional peptide modifications, which were not covered by the database or could not even be identified by the open modification search approach. Furthermore, peptides containing unexpected semi-tryptic or nonspecific cleavages may also add to this fraction [21].

**Figure 7:**
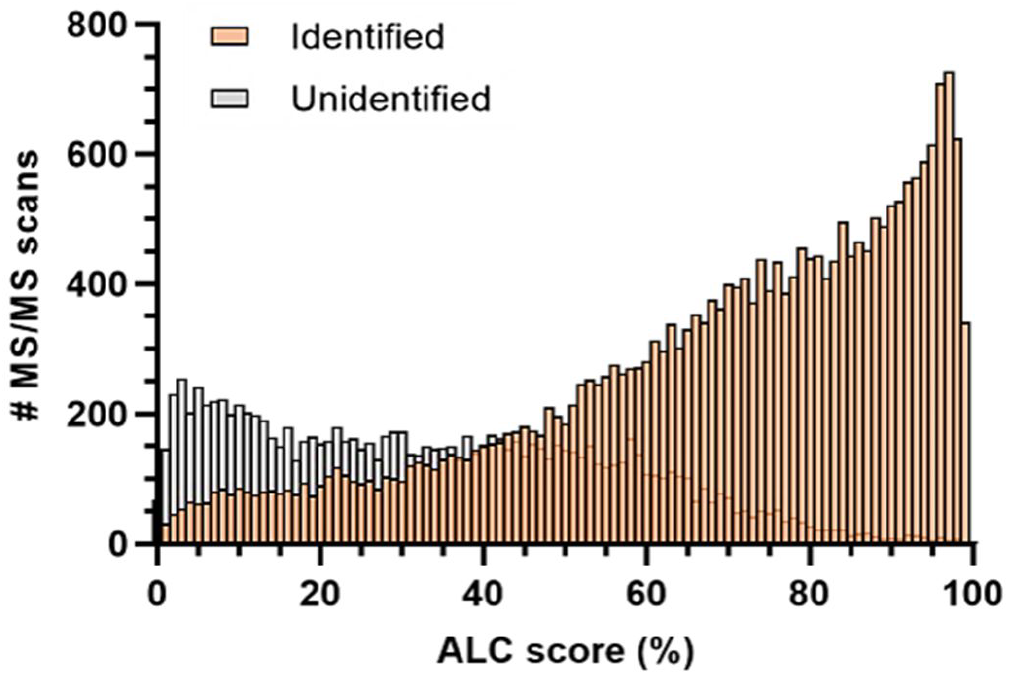
Assessing the spectral quality of the acquired fragmentation spectra. The Average Local Confidence (ALC) score distribution of identified (orange) and unidentified (grey) spectra, as shown for protocol 13 (biological replicate 1). In general, peptides with higher ALC score are more likely to provide a confident match during database search (because of the better fragment ion coverage of the peptide backbone, and the possibly ‘cleaner’ spectrum). The unidentified fraction from protocol 13 (grey) shows an ACL score distribution at predominantly lower numbers, indicating that the unidentified spectra largely derive from very low quality spectra, insufficient to provide a confident peptide sequence match. The fraction of higher ALC scoring spectra presumably derive from sequence variants or unexpected modifications not present in the database. Taxonomic impurities or sample carry over can be excluded.

## CONCLUSION

Our systematic study on the unicellular organism yeast demonstrates the strong impact of several elements within sample preparation protocols, on various aspects of the proteomics analysis outcome. The total number of obtained peptide spectrum matches ranged from approx. 1×10^4^ to 3×10^4^, the proteome coverage ranged from below 15% to approx. 30%, and the overall matched spectra varied in the range from approx. 50 to 70%. The depth of the proteome coverage was heavily affected by the sample quenching protocol, where cells arrested by TCA resulted in a higher number of protein identifications compared to methanol-arrested cells. Furthermore, filter-aided procedures, when combined with Urea, outperformed other protocols in terms of the number of protein identifications.

The use of unrestricted modification search moreover enabled to examine for reagent-induced peptide modifications. The overall frequency of reagent-specific modifications was for some protocols significantly lower (particularly for protocols 8 and 14), albeit that also correlated with the lower number of spectra and decreased total ion current (TIC) observed for those protocols. Our analysis confirmed previously identified sample processing induced modifications, but also revealed unexpected modifications, such as those related to acetone, alkylating agents as well as filter materials.

Approximately 70% of the overall acquired MS/MS spectra could be identified when using filter-aided as well as the best performing in-solution protocols. The unidentified spectra were predominantly of low-quality, lacking sufficient peptide fragment coverage for a confident identification. A maximum proteome coverage and reduced number of reagent-induced modifications were obtained for the filter-aided approach, even in the presence of Urea buffer (when using moderate incubation temperatures). However, this approach appeared to be more laborious and requires the availability of suitable filters and centrifuges. A comparable performance was obtained with the best performing in-solution protocols, such as 7 and 11. More specialised protocols, such as S-trap [81] and ultrasonic FASP digestion [82] were not evaluated, but are expected to enable a similarly good proteome coverage at reduced processing times.

Ultimately, we systematically evaluated a matrix of sample preparation protocols for the unicellular model eukaryote yeast, and provide a large resource of protocols and associated mass spectrometric raw data. These enable the selection of suitable sample preparation elements, the design of more specialised procedures and support the evaluation and analysis of (native) peptide modifications in yeast.

## AUTHOR INFORMATION

### Author Contributions

MP, PDL and MDR designed the experiments. MDR, EK and WB performed experiments. MDR and MP analyzed the data. MDR and MP wrote the manuscript. All authors have given approval to the final version of the manuscript.

### Notes

This work was supported by a TU Delft startup fund. Conflict of interest—The authors declare no competing interests.

## ACKNOWLEDGMENT

The authors are grateful to valuable discussions with their colleagues from the department of Biotechnology and acknowledge Carol de Ram for technical support.

