## Supplementary material for "A systematic evaluation of yeast sample preparation protocols for spectral identifications, proteome coverage and post-isolation modifications": SI document

### SI information material to:

Maxime den Ridder, Ewout Knibbe, Wiebeke van den Brandeler, Pascale Daran-Lapujade and Martin Pabst*
Delft University of Technology, Department of Biotechnology, van der Maasweg 9, 2629 HZ Delft, The Netherlands

**Table S1:** Growth characteristics of IMX372 (MG strain) in glucose-limited aerobic chemostat cultures at a dilution rate of 0.10 $h^{-1}$.

| Bioreactor | Replicate 1 | Replicate 2 |
|---|---|---|
| Biomass yield ($g_x$ $g_{sucrose}^{-1}$) | 0.46 | 0.44 |
| $q_{glucose}$ (mmol $g_x^{-1}$ $h^{-1}$) | -1.211 | -1.264 |
| $q_{CO2}$ (mmol $g_x^{-1}$ $h^{-1}$) | 3.15 | 3.24 |
| $q_{O2}$ (mmol $g_x^{-1}$ $h^{-1}$) | -3.10 | -3.22 |
| Respiratory quotient | 1.02 | 1.00 |
| Carbon recovery (%) | 100.01 | 97.09 |
| Actual dilution rate ($h^{-1}$) | 0.101 | 0.101 |

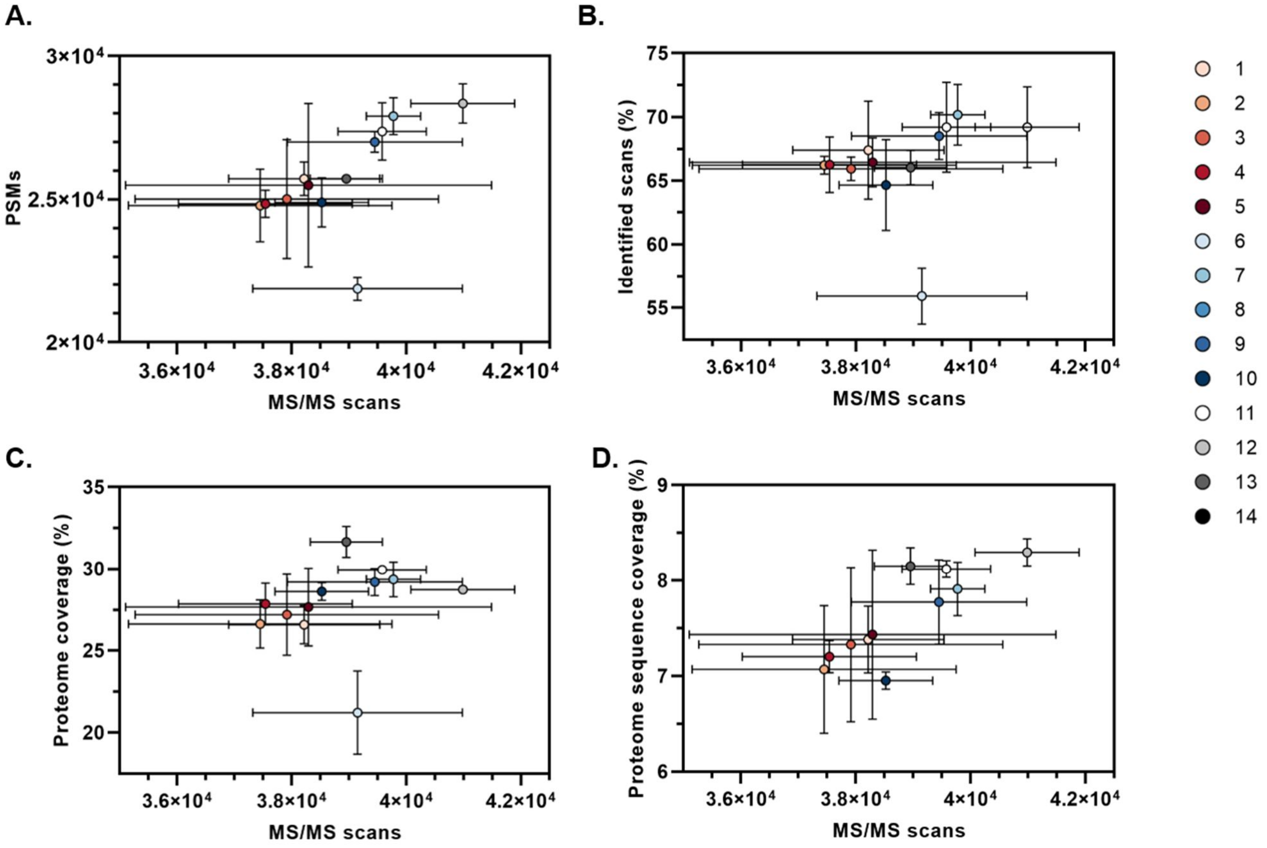


**Figure S1: Achieved sequence identification rates, proteome and sequence coverage for the different sample preparation protocols. (zoomed version of figure 2).** The number of Peptide Spectrum Matches (PSMs) (**A**), identified scans (%) (**B**), proteome coverage (%) (**C**) and proteome sequence coverage (**D**) were plotted against the number of MS/MS scans obtained per protocol. Each colored circle represents the average of four technical replicates obtained from two biological replicates, while the standard deviation is represented by the error bars. The proteome coverage was calculated based on the proteins identified per protocol as a percentage of the total number of proteins in yeast (known ORFs). In addition, the proteome sequence coverage was calculated based on the identified amino acids in the proteomics experiments, as a percentage of the total proteome amino acid sequence (sequence coverage).

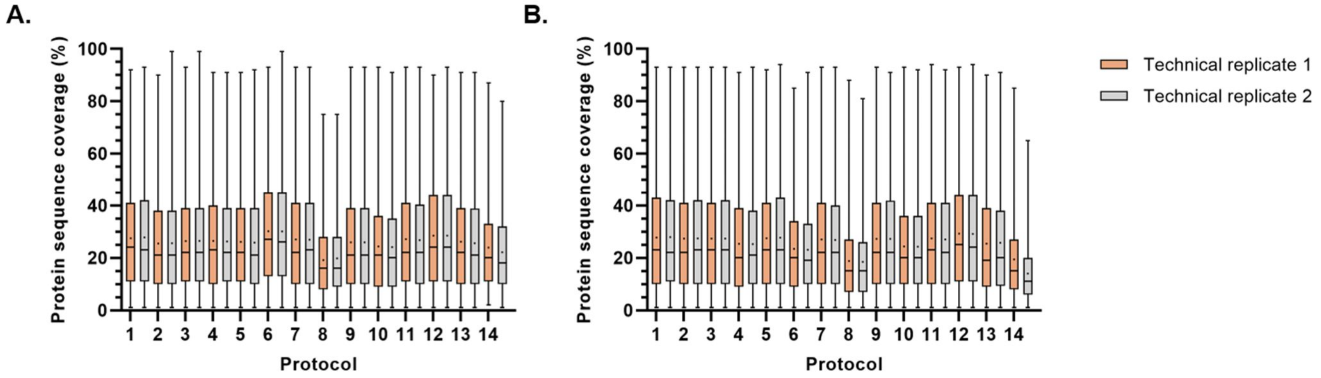


**Figure S2: Boxplot distribution of protein sequence coverage obtained after applying different sample preparation protocols (1–14).** MS/MS spectra were identified using conventional database matching with PEAKS-DB in which at least 2 unique peptides had to be found for a reliable protein identification. The distribution of the sequence coverage of these identified proteins is shown here for each sample for both technical replicates (orange and gray). The average sequence coverage (·) and median (-) are shown in the boxplots. The average sequence coverage across all samples was 26 % and 25 %, for biological replicate 1 (**A**) and 2 (**B**), respectively.

### Detailed description of employed yeast whole cell lysate proteomics sample preparation protocols

| # | Sampling method | Lysis buffer | Reducing agent | Alkylation agent | Protein precipitation | SPE elution buffer |
|---|---|---|---|---|---|---|
| **1** | TCA | SDS | DTT | IAA | Acetone | **TFA** |
| **2** | TCA | SDS | DTT | IAA | Acetone | **FA + ABC** |
| **3** | TCA | SDS | DTT | IAA | Acetone | **ABC** |
| **4** | TCA | SDS | DTT | IAA | Acetone | **MeOH** |
| **5** | TCA | SDS | DTT | IAA | Acetone | **FA** |
| **6** | **MeOH** | SDS | DTT | IAA | Acetone | TFA + ABC |
| **7** | TCA | SDS | DTT | IAA | Acetone | TFA + ABC |
| **8** | TCA | SDS | DTT | IAA | **TCA** | TFA + ABC |
| **9** | TCA | SDS | DTT | **AA** | Acetone | TFA + ABC |
| **10** | TCA | SDS | DTT | **-** | Acetone | TFA + ABC |
| **11** | TCA | SDS | **TCEP** | IAA | Acetone | TFA + ABC |
| **12** | TCA | **UREA** | DTT | IAA | Acetone | TFA + ABC |
| **13** | TCA | **UREA** | DTT | IAA | **FASP** | TFA + ABC |
| **14** | TCA | **SDS** | DTT | IAA | **FASP** | TFA + ABC |

#### ADDITIONAL INFORMATION MATERIALS

- Trichloroacetic acid (TCA, Merck Sigma, Cat. No. T0699)
- Methanol (MeOH, Thermo Fisher, Cat. No. 15654570)
- Triethylammonium bicarbonate (TEAB, Merck Sigma, Cat. No. T7408)
- Sodium dodecyl sulfate (SDS, Merck Sigma, Cat. No. L4522)
- Urea (Merck Sigma, Cat. No. U5378)
- Phosphatase inhibitor (Merck Sigma, Cat. No. 524625)
- Protease inhibitor (Merck Sigma, Cat. No. P8215)
- Acid washed glass beads (Merk Sigma, Cat. No. G8772)
- Shock resistant tubes (Labservices BV, Cat. No. 330TX)
- LoBind tubes, 1.5 mL (Eppendorf Cat. No. 0030108434)
- Dithiothreitol (DTT, Merck Sigma, Cat. No. 43815)
- Tris(2-carboxyethyl)phosphine (TCEP, Merck Sigma, Cat. No. C4706)
- Iodoacetamide (IAA, Merck Sigma, Cat. No. I1149)
- Acrylamide (AA, Merck Sigma, Cat. No. A9099)
- Acetone (Merck Sigma, Cat. No. 650501)
- Sample filters (Merck-Millipore, Microcon 10 kDa, Cat. No. MRCPRT010)
- Trypsin (Promega, Cat. No. V5111)
- Ammonium bicarbonate (ABC, Merck Sigma, Cat. No. 09830)
- Acetonitrile (ACN, Thermo Fisher, Cat. No. 10489553)
- Formic acid (FA, Thermo Fisher, Cat. No. 10596814)
- Trifluoroacetic acid (TFA, Merck Sigma, Cat. No. 302031)

#### DETAILED DESCRIPTION PROTOCOL 1

1. **Biomass collection and quenching.** Yeast biomass (1 mL, at 2.6 g/L dry weight) was collected from steady state cultures. The samples were collected in multifold in trichloroacetic acid (TCA) with a final concentration of 10%**.** Samples were centrifuged using a bench top centrifuge (Eppendorf) at 4000 g for 5 min at 4°C. Cell pellets were frozen at -80°C until further processed.

2. **Cell lysis.** Yeast cell culture pellets were resuspended in lysis buffer composed of 100 mM triethylamonium bicarbonate (TEAB) containing 1% SDS and phosphatase/protease inhibitors. Yeast cells were lysed by glass beads beating using a Mini-Beadbeater-24. Samples were shaken in the presence of 0.1 g acid washed glass beads using shock resistant tubes 10 times for 1 min with 1 min rest on ice in-between.

3. **In-solution digestion.** The protein supernatant was separated from insoluble cell debris by centrifugation at 14.000 rpm for 5 min at 10°C using a benchtop centrifuge (Eppendorf). Proteins were reduced by addition of 5 mM Dithiothreitol (DTT) (final concentration) for 1 hour at 37°C using an Eppendorf ThermoMixer. Subsequently, the proteins were alkylated in the dark by addition of 15 mM iodoacetamide for 30 min at room temperature. Protein precipitation was performed by addition of four volumes of ice-cold acetone (-20°C) and proceeded for 1 hour at -20°C. The solution was centrifuged at 14.000 rpm using a bench-top centrifuge at 10°C to collect the cell pellet. The acetone supernatant was discarded. Proteins were then solubilized using 100 mM ammonium bicarbonate (ABC) and proteolytic digestion was performed by addition of Trypsin (Promega, Madison, WI), to a 1:100 enzyme to protein ratio (v/v), and incubated at 37°C overnight.

4. **Solid phase extraction of the proteolytic digest.** Solid phase extraction of the proteolytic digest was performed with an Oasis HLB 96-well µElution plate (Waters, Milford, USA). The columns were first washed with 750 µL methanol and equilibrated with 1000 µL water. Samples were then loaded and washed twice with 5% MeOH. Peptide fractions were eluted in twice using <u>80% MeOH buffers containing 0.01% trifluoroacetic acid (TFA)</u>. Eluates were collected in 1.5 mL LoBind tubes (Eppendorf) and dried using a SpeedVac concentrator for two hours (temperature was set to 50°C for one hour, and to RT for the second hour). Dried peptides were resuspended in 3% acetonitrile (ACN) / 0.01% TFA prior to MS-analysis to give an approximate concentration of 500 ng per µL.

#### DETAILED DESCRIPTION PROTOCOL 2

1. **Sample biomass collection.** Yeast biomass (1 mL) was collected from steady state cultures. The samples were collected in multifold in trichloroacetic acid (TCA) with a final concentration of 10%**.** Samples were centrifuged using a bench top centrifuge (Eppendorf) at 4000 g for 5 min at 4°C. Cell pellets were frozen at -80°C.

2. **Cell lysis.** Yeast cell culture pellets were resuspended in lysis buffer composed of 100 mM triethylamonium bicarbonate (TEAB) containing 1% SDS and phosphatase/protease inhibitors. Yeast cells were lysed by glass beads beating using a Mini-Beadbeater-24. Samples were shaken in the presence of 0.1 g acid washed glass beads using shock resistant tubes 10 times for 1 min with 1 min rest on ice in-between.

3. **In-solution digestion.** The protein supernatant was separated from insoluble cell debris by centrifugation at 14.000 rpm for 5 min at 10°C using a benchtop centrifuge (Eppendorf). Proteins were reduced by addition of 5 mM

Dithiothreitol (DTT) (final concentration) for 1 hour at 37°C using an Eppendorf ThermoMixer. Subsequently, the proteins were alkylated in the dark by addition of 15 mM iodoacetamide for 30 min at room temperature. Protein precipitation was performed by addition of four volumes of ice-cold acetone (-20° C) and proceeded for 1 hour at -20°C. The solution was centrifuged at 14.000 rpm using a bench-top centrifuge at 10°C to collect the cell pellet. The acetone supernatant was discarded. Proteins were then solubilized using 100 mM ammonium bicarbonate (ABC) and proteolytic digestion was performed by addition of Trypsin (Promega, Madison, WI), to a 1:100 enzyme to protein ratio (v/v), and incubated at 37°C overnight.

4. **Solid phase extraction of the proteolytic digest.** Solid phase extraction of the proteolytic digest was performed with an Oasis HLB 96-well µElution plate (Waters, Milford, USA). The columns were first washed with 750 µL methanol and equilibrated with 1000 µL water. Samples were then loaded and washed twice with 5% MeOH. Peptide fractions were eluted in two 80% MeOH buffers containing 0.1% Formic Acid (FA) and 1 mM ABC, separately. Eluates were collected in 1.5 mL LoBind tubes (Eppendorf) and dried using a SpeedVac concentrator for two hours (temperature was set to 50°C for one hour, and to RT for the second hour). Dried peptides were resuspended in 3% acetonitrile (ACN) / 0.01% TFA prior to MS-analysis to give an approximate concentration of 500 ng per µL.

**DETAILED DESCRIPTION PROTOCOL 3**

1. **Sample biomass collection.** Yeast biomass (1 mL) was collected from steady state cultures. The samples were collected in multifold in trichloroacetic acid (TCA) with a final concentration of 10%. Samples were centrifuged using a bench top centrifuge (Eppendorf) at 4000 g for 5 min at 4°C. Cell pellets were frozen at -80°C.

2. **Cell lysis.** Yeast cell culture pellets were resuspended in lysis buffer composed of 100 mM triethylamonium bicarbonate (TEAB) containing 1% SDS and phosphatase/protease inhibitors. Yeast cells were lysed by glass beads beating using a Mini-Beadbeater-24. Samples were shaken in the presence of 0.1 g acid washed glass beads using shock resistant tubes 10 times for 1 min with 1 min rest on ice in-between.

3. **In-solution digestion.** The protein supernatant was separated from insoluble cell debris by centrifugation at 14.000 rpm for 5 min at 10°C using a benchtop centrifuge (Eppendorf). Proteins were reduced by addition of 5 mM Dithiothreitol (DTT) (final concentration) for 1 hour at 37°C using an Eppendorf ThermoMixer. Subsequently, the proteins were alkylated in the dark by addition of 15 mM iodoacetamide for 30 min at room temperature. Protein precipitation was performed by addition of four volumes of ice-cold acetone (-20° C) and proceeded for 1 hour at -20°C. The solution was centrifuged at 14.000 rpm using a bench-top centrifuge at 10°C to collect the cell pellet. The acetone supernatant was discarded. Proteins were then solubilized using 100 mM ammonium bicarbonate (ABC) and proteolytic digestion was performed by addition of Trypsin (Promega, Madison, WI), to a 1:100 enzyme to protein ratio (v/v), and incubated at 37°C overnight.

4. **Solid phase extraction of the proteolytic digest.** Solid phase extraction of the proteolytic digest was performed with an Oasis HLB 96-well µElution plate (Waters, Milford, USA). The columns were first washed with 750 µL methanol and equilibrated with 1000 µL water. Samples were then loaded and washed twice with 5% MeOH. Peptide fractions were eluted in twice using 80% MeOH buffers containing 1 mM ABC. Eluates were collected in 1.5 mL LoBind tubes (Eppendorf) and dried using a SpeedVac concentrator for two hours (temperature was set to 50°C for one hour, and to

RT for the second hour). Dried peptides were resuspended in 3% acetonitrile (ACN) / 0.01% TFA prior to MS-analysis to give an approximate concentration of 500 ng per µL.

**DETAILED DESCRIPTION PROTOCOL 4**

1. **Sample biomass collection.** Yeast biomass (1 mL) was collected from steady state cultures. The samples were collected in multifold in trichloroacetic acid (TCA) with a final concentration of 10%**.** Samples were centrifuged using a bench top centrifuge (Eppendorf) at 4000 g for 5 min at 4°C. Cell pellets were frozen at -80°C.

2. **Cell lysis.** Yeast cell culture pellets were resuspended in lysis buffer composed of 100 mM triethylamonium bicarbonate (TEAB) containing 1% SDS and phosphatase/protease inhibitors. Yeast cells were lysed by glass beads beating using a Mini-Beadbeater-24. Samples were shaken in the presence of 0.1 g acid washed glass beads using shock resistant tubes 10 times for 1 min with 1 min rest on ice in-between.

3. **In-solution digestion.** The protein supernatant was separated from insoluble cell debris by centrifugation at 14.000 rpm for 5 min at 10°C using a benchtop centrifuge (Eppendorf). Proteins were reduced by addition of 5 mM Dithiothreitol (DTT) (final concentration) for 1 hour at 37°C using an Eppendorf ThermoMixer. Subsequently, the proteins were alkylated in the dark by addition of 15 mM iodoacetamide for 30 min at room temperature. Protein precipitation was performed by addition of four volumes of ice-cold acetone (-20° C) and proceeded for 1 hour at -20°C. The solution was centrifuged at 14.000 rpm using a bench-top centrifuge at 10°C to collect the cell pellet. The acetone supernatant was discarded. Proteins were then solubilized using 100 mM ammonium bicarbonate (ABC) and proteolytic digestion was performed by addition of Trypsin (Promega, Madison, WI), to a 1:100 enzyme to protein (v/v), and incubated at 37°C overnight.

4. **Solid phase extraction of the proteolytic digest.** Solid phase extraction of the proteolytic digest was performed with an Oasis HLB 96-well µElution plate (Waters, Milford, USA). The columns were first washed with 750 µL methanol and equilibrated with 1000 µL water. Samples were then loaded and washed twice with 5% MeOH. Peptide fractions were eluted in twice using <u>80% MeOH buffers containing 1 mM ABC</u>. Eluates were collected in 1.5 mL LoBind tubes (Eppendorf) and dried using a SpeedVac concentrator for two hours (temperature was set to 50°C for one hour, and to RT for the second hour). Dried peptides were resuspended in 3% acetonitrile (ACN) / 0.01% TFA prior to MS-analysis to give an approximate concentration of 500 ng per µL.

**DETAILED DESCRIPTION PROTOCOL 5**

1. **Sample biomass collection.** Yeast biomass (1 mL) was collected from steady state cultures. The samples were collected in multifold in trichloroacetic acid (TCA) with a final concentration of 10%. Samples were centrifuged using a bench top centrifuge (Eppendorf) at 4000 g for 5 min at 4°C. Cell pellets were frozen at -80°C.

2. **Cell lysis.** Yeast cell culture pellets were resuspended in lysis buffer composed of 100 mM triethylamonium bicarbonate (TEAB) containing 1% SDS and phosphatase/protease inhibitors. Yeast cells were lysed by glass beads beating using a Mini-Beadbeater-24. Samples were shaken in the presence of 0.1 g acid washed glass beads using shock resistant tubes 10 times for 1 min with 1 min rest on ice in-between.

3. **In-solution digestion.** The protein supernatant was separated from insoluble cell debris by centrifugation at 14.000 rpm for 5 min at 10°C using a benchtop centrifuge (Eppendorf). Proteins were reduced by addition of 5 mM Dithiothreitol (DTT) (final concentration) for 1 hour at 37°C using an Eppendorf ThermoMixer. Subsequently, the proteins were alkylated in the dark by addition of 15 mM iodoacetamide for 30 min at room temperature. Protein precipitation was performed by addition of four volumes of ice-cold acetone (-20° C) and proceeded for 1 hour at -20°C. The solution was centrifuged at 14.000 rpm using a bench-top centrifuge at 10°C to collect the cell pellet. The acetone supernatant was discarded. Proteins were then solubilized using 100 mM ammonium bicarbonate (ABC) and proteolytic digestion was performed by addition of Trypsin (Promega, Madison, WI), to a 1:100 enzyme to protein ratio (v/v), and incubated at 37°C overnight.

4. **Solid phase extraction of the proteolytic digest.** Solid phase extraction of the proteolytic digest was performed with an Oasis HLB 96-well µElution plate (Waters, Milford, USA). The columns were first washed with 750 µL methanol and equilibrated with 1000 µL water. Samples were then loaded and washed twice with 5% MeOH. Peptide fractions were eluted in twice using <u>80% MeOH buffers containing 0.1% FA</u>. Eluates were collected in 1.5 mL LoBind tubes (Eppendorf) and dried using a SpeedVac concentrator for two hours (temperature was set to 50°C for one hour, and to RT for the second hour). Dried peptides were resuspended in 3% acetonitrile (ACN) / 0.01% TFA prior to MS-analysis to give an approximate concentration of 500 ng per µL.

**DETAILED DESCRIPTION PROTOCOL 6**

1. **Sample biomass collection.** Yeast biomass (1 mL) was collected from steady state cultures. The samples were collected in multifold <u>in five volumes of ice-cold methanol (MeOH).</u> Samples were centrifuged using a bench top centrifuge (Eppendorf) at 4000 g for 5 min at 4°C. Cell pellets were frozen at -80°C.

2. **Cell lysis.** Yeast cell culture pellets were resuspended in lysis buffer composed of 100 mM triethylamonium bicarbonate (TEAB) containing 1% SDS and phosphatase/protease inhibitors. Yeast cells were lysed by glass beads beating using a Mini-Beadbeater-24. Samples were shaken in the presence of 0.1 g acid washed glass beads using shock resistant tubes 10 times for 1 min with 1 min rest on ice in-between.

3. **In-solution digestion.** The protein supernatant was separated from insoluble cell debris by centrifugation at 14.000 rpm for 5 min at 10°C using a benchtop centrifuge (Eppendorf). Proteins were reduced by addition of 5 mM Dithiothreitol (DTT) (final concentration) for 1 hour at 37°C using an Eppendorf ThermoMixer. Subsequently, the proteins were alkylated in the dark by addition of 15 mM iodoacetamide for 30 min at room temperature. Protein precipitation was performed by addition of four volumes of ice-cold acetone (-20° C) and proceeded for 1 hour at -20°C. The solution was centrifuged at 14.000 rpm using a bench-top centrifuge at 10°C to collect the cell pellet. The acetone supernatant was discarded. Proteins were then solubilized using 100 mM ammonium bicarbonate (ABC) and proteolytic digestion was performed by addition of Trypsin (Promega, Madison, WI), to a 1:100 enzyme to protein ratio (v/v), and incubated at 37°C overnight.

**4.** **Solid phase extraction of the proteolytic digest.** Solid phase extraction of the proteolytic digest was performed with an Oasis HLB 96-well µElution plate (Waters, Milford, USA). The columns were first washed with 750 µL methanol and equilibrated with 1000 µL water. Samples were then loaded and washed twice with 5% MeOH. Peptide fractions were eluted in two 80% MeOH buffers containing 0.01% TFA and 1 mM ABC, separately. Eluates were collected in 1.5 mL LoBind tubes (Eppendorf) and dried using a SpeedVac concentrator for two hours (temperature was set to 50°C for one hour, and to RT for the second hour). Dried peptides were resuspended in 3% acetonitrile (ACN) / 0.01% TFA prior to MS-analysis to give an approximate concentration of 500 ng per µL.

**DETAILED DESCRIPTION PROTOCOL 7**

1. **Sample biomass collection.** Yeast biomass (1 mL) was collected from steady state cultures. The samples were collected in multifold in trichloroacetic acid (TCA) with a final concentration of 10%. Samples were centrifuged using a bench top centrifuge (Eppendorf) at 4000 g for 5 min at 4°C. Cell pellets were frozen at -80°C.

2. **Cell lysis.** Yeast cell culture pellets were resuspended in lysis buffer composed of 100 mM triethylamonium bicarbonate (TEAB) containing 1% SDS and phosphatase/protease inhibitors. Yeast cells were lysed by glass beads beating using a Mini-Beadbeater-24. Samples were shaken in the presence of 0.1 g acid washed glass beads using shock resistant tubes 10 times for 1 min with 1 min rest on ice in-between.

3. **In-solution digestion.** The protein supernatant was separated from insoluble cell debris by centrifugation at 14.000 rpm for 5 min at 10°C using a benchtop centrifuge (Eppendorf). Proteins were reduced by addition of 5 mM Dithiothreitol (DTT) (final concentration) for 1 hour at 37°C using an Eppendorf ThermoMixer. Subsequently, the proteins were alkylated in the dark by addition of 15 mM iodoacetamide for 30 min at room temperature. Protein precipitation was performed by addition of four volumes of ice-cold acetone (-20° C) and proceeded for 1 hour at -20°C. The solution was centrifuged at 14.000 rpm using a bench-top centrifuge at 10°C to collect the cell pellet. The acetone supernatant was discarded. Proteins were then solubilized using 100 mM ammonium bicarbonate (ABC) and proteolytic digestion was performed by addition of Trypsin (Promega, Madison, WI), to a 1:100 enzyme to protein ratio (v/v), and incubated at 37°C overnight.

4. **Solid phase extraction of the proteolytic digest.** Solid phase extraction of the proteolytic digest was performed with an Oasis HLB 96-well µElution plate (Waters, Milford, USA). The columns were first washed with 750 µL methanol and equilibrated with 1000 µL water. Samples were then loaded and washed twice with 5% MeOH. Peptide fractions were eluted in two 80% MeOH buffers containing 0.01% TFA and 1 mM ABC, separately. Eluates were collected in 1.5 mL LoBind tubes (Eppendorf) and dried using a SpeedVac concentrator for two hours (temperature was set to 50°C for one hour, and to RT for the second hour). Dried peptides were resuspended in 3% acetonitrile (ACN) / 0.01% TFA prior to MS-analysis to give an approximate concentration of 500 ng per µL.

**DETAILED DESCRIPTION PROTOCOL 8**

1. **Sample biomass collection.** Yeast biomass (1 mL) was collected from steady state cultures. The samples were collected in multifold in trichloroacetic acid (TCA) with a final concentration of 10%. Samples were centrifuged using a bench top centrifuge (Eppendorf) at 4000 g for 5 min at 4°C. Cell pellets were frozen at -80°C.

2. **Cell lysis.** Yeast cell culture pellets were resuspended in lysis buffer composed of 100 mM triethylamonium bicarbonate (TEAB) containing 1% SDS and phosphatase/protease inhibitors. Yeast cells were lysed by glass beads beating using a

Mini-Beadbeater-24. Samples were shaken in the presence of 0.1 g acid washed glass beads using shock resistant tubes 10 times for 1 min with 1 min rest on ice in-between.

3. **In-solution digestion.** The protein supernatant was separated from insoluble cell debris by centrifugation at 14.000 rpm for 5 min at 10°C using a benchtop centrifuge (Eppendorf). Proteins were reduced by addition of 5 mM Dithiothreitol (DTT) (final concentration) for 1 hour at 37°C using an Eppendorf ThermoMixer. Subsequently, the proteins were alkylated in the dark by addition of 15 mM iodoacetamide for 30 min at room temperature. Protein precipitation was performed by addition of TCA to a final concentration of 20% and proceeded for 30 min at 4°C. The solution was centrifuged at 14.000 rpm using a bench-top centrifuge at 10°C to collect the cell pellet. The acetone supernatant was discarded. Proteins were then solubilized using 100 mM ammonium bicarbonate (ABC) and proteolytic digestion was performed by addition of Trypsin (Promega, Madison, WI), to a 1:100 enzyme to protein ratio (v/v), and incubated at 37°C overnight.

4. **Solid phase extraction of the proteolytic digest.** Solid phase extraction of the proteolytic digest was performed with an Oasis HLB 96-well µElution plate (Waters, Milford, USA). The columns were first washed with 750 µL methanol and equilibrated with 1000 µL water. Samples were then loaded and washed twice with 5% MeOH. Peptide fractions were eluted in two 80% MeOH buffers containing 0.01% TFA and 1 mM ABC, separately. Eluates were collected in 1.5 mL LoBind tubes (Eppendorf) and dried using a SpeedVac concentrator for two hours (temperature was set to 50°C for one hour, and to RT for the second hour). Dried peptides were resuspended in 3% acetonitrile (ACN) / 0.01% TFA prior to MS-analysis to give an approximate concentration of 500 ng per µL.

**DETAILED DESCRIPTION PROTOCOL 9**

1. **Sample biomass collection.** Yeast biomass (1 mL) was collected from steady state cultures. The samples were collected in multifold in trichloroacetic acid (TCA) with a final concentration of 10%. Samples were centrifuged using a bench top centrifuge (Eppendorf) at 4000g for 5 min at 4°C. Cell pellets were frozen at -80°C.

2. **Cell lysis.** Yeast cell culture pellets were resuspended in lysis buffer composed of 100 mM triethylamonium bicarbonate (TEAB) containing 1% SDS and phosphatase/protease inhibitors. Yeast cells were lysed by glass beads beating using a Mini-Beadbeater-24. Samples were shaken in the presence of 0.1 g acid washed glass beads using shock resistant tubes 10 times for 1 min with 1 min rest on ice in-between.

3. **In-solution digestion.** The protein supernatant was separated from insoluble cell debris by centrifugation at 14.000 rpm for 5 min at 10°C using a benchtop centrifuge (Eppendorf). Proteins were reduced by addition of 5 mM Dithiothreitol (DTT) (final concentration) for 1 hour at 37°C using an Eppendorf ThermoMixer. Subsequently, the proteins were alkylated in the dark by addition of 50 mM acrylamide for 1 hour at room temperature. Protein precipitation was performed by addition of four volumes of ice-cold acetone (-20°C) and proceeded for 1 hour at -20°C. The solution was centrifuged at 14.000 rpm using a bench-top centrifuge at 10°C to collect the cell pellet. The acetone supernatant was discarded. Proteins were then solubilized using 100 mM ammonium bicarbonate (ABC) and proteolytic digestion was performed by addition of Trypsin (Promega, Madison, WI), to a 1:100 enzyme to protein ratio (v/v), and incubated at 37°C overnight.

4. **Solid phase extraction of the proteolytic digest.** Solid phase extraction of the proteolytic digest was performed with an Oasis HLB 96-well μElution plate (Waters, Milford, USA). The columns were first washed with 750 μL methanol and equilibrated with 1000 μL water. Samples were then loaded and washed twice with 5% MeOH. Peptide fractions were eluted in two 80% MeOH buffers containing 0.01% TFA and 1 mM ABC, separately. Eluates were collected in 1.5 mL LoBind tubes (Eppendorf) and dried using a SpeedVac concentrator for two hours (temperature was set to 50°C for one hour, and to RT for the second hour). Dried peptides were resuspended in 3% acetonitrile (ACN) / 0.01% TFA prior to MS-analysis to give an approximate concentration of 500 ng per μL.

**DETAILED DESCRIPTION PROTOCOL 10**

1. **Sample biomass collection.** Yeast biomass (1 mL) was collected from steady state cultures. The samples were collected in multifold in trichloroacetic acid (TCA) with a final concentration of 10%. Samples were centrifuged using a bench top centrifuge (Eppendorf) at 4000 g for 5 min at 4°C. Cell pellets were frozen at -80°C.

2. **Cell lysis.** Yeast cell culture pellets were resuspended in lysis buffer composed of 100 mM triethylamonium bicarbonate (TEAB) containing 1% SDS and phosphatase/protease inhibitors. Yeast cells were lysed by glass beads beating using a Mini-Beadbeater-24. Samples were shaken in the presence of 0.1 g acid washed glass beads using shock resistant tubes 10 times for 1 min with 1 min rest on ice in-between.

3. **In-solution digestion.** The protein supernatant was separated from insoluble cell debris by centrifugation at 14.000 rpm for 5 min at 10°C using a benchtop centrifuge (Eppendorf). Proteins were reduced by addition of 5 mM Dithiothreitol (DTT) (final concentration) for 1 hour at 37°C. The proteins were not alkylated. Protein precipitation was performed by addition of four volumes of ice-cold acetone (-20°C) and proceeded for 1 hour at -20°C. The solution was centrifuged at 14.000 rpm using a bench-top centrifuge at 10°C to collect the cell pellet. The acetone supernatant was discarded. Proteins were then solubilized using 100 mM ammonium bicarbonate (ABC) and proteolytic digestion was performed by addition of Trypsin (Promega, Madison, WI), to a 1:100 enzyme to protein ratio (v/v), and incubated at 37°C overnight.

4. **Solid phase extraction of the proteolytic digest.** Solid phase extraction of the proteolytic digest was performed with an Oasis HLB 96-well μElution plate (Waters, Milford, USA). The columns were first washed with 750 μL methanol and equilibrated with 1000 μL water. Samples were then loaded and washed twice with 5% MeOH. Peptide fractions were eluted in two 80% MeOH buffers containing 0.01% TFA and 1 mM ABC, separately. Eluates were collected in 1.5 mL LoBind tubes (Eppendorf) and dried using a SpeedVac concentrator for two hours (temperature was set to 50°C for one hour, and to RT for the second hour). Dried peptides were resuspended in 3% acetonitrile (ACN) / 0.01% TFA prior to MS-analysis to give an approximate concentration of 500 ng per μL.

**DETAILED DESCRIPTION PROTOCOL 11**

1. **Sample biomass collection.** Yeast biomass (1 mL) was collected from steady state cultures. The samples were collected in multifold in trichloroacetic acid (TCA) with a final concentration of 10%. Samples were centrifuged using a bench top centrifuge (Eppendorf) at 4000 g for 5 min at 4°C. Cell pellets were frozen at -80°C.

2. **Cell lysis.** Yeast cell culture pellets were resuspended in lysis buffer composed of 100 mM triethylamonium bicarbonate (TEAB) containing 1% SDS and phosphatase/protease inhibitors. Yeast cells were lysed by glass beads beating using a Mini-Beadbeater-24. Samples were shaken in the presence of 0.1 g acid washed glass beads using shock resistant tubes 10 times for 1 min with 1 min rest on ice in-between.

3. **In-solution digestion.** The protein supernatant was separated from insoluble cell debris by centrifugation at 14.000 rpm for 5 min at 10°C using a benchtop centrifuge (Eppendorf). Proteins were reduced by addition of 5 mM TCEP (final concentration) for 30 min at 56°C using an Eppendorf ThermoMixer. Subsequently, the proteins were alkylated in the dark by addition of 15 mM iodoacetamide for 30 min at room temperature. Protein precipitation was performed by addition of four volumes of ice-cold acetone (-20°C) and proceeded for 1 hour at -20°C. The solution was centrifuged at 14.000 rpm using a bench-top centrifuge at 10°C to collect the cell pellet. The acetone supernatant was discarded. Proteins were then solubilized using 100 mM ammonium bicarbonate (ABC) and proteolytic digestion was performed by addition of Trypsin (Promega, Madison, WI), to a 1:100 enzyme to protein ratio (v/v), and incubated at 37°C overnight.

4. **Solid phase extraction of the proteolytic digest.** Solid phase extraction of the proteolytic digest was performed with an Oasis HLB 96-well µElution plate (Waters, Milford, USA). The columns were first washed with 750 µL methanol and equilibrated with 1000 µL water. Samples were then loaded and washed twice with 5% MeOH. Peptide fractions were eluted in two 80% MeOH buffers containing 0.01% TFA and 1 mM ABC, separately. Eluates were collected in 1.5 mL LoBind tubes (Eppendorf) and dried using a SpeedVac concentrator for two hours (temperature was set to 50°C for one hour, and to RT for the second hour). Dried peptides were resuspended in 3% acetonitrile (ACN) / 0.01% TFA prior to MS-analysis to give an approximate concentration of 500 ng per µL.

**DETAILED DESCRIPTION PROTOCOL 12**

1. **Sample biomass collection.** Yeast biomass (1 mL) was collected from steady state cultures. The samples were collected in multifold in trichloroacetic acid (TCA) with a final concentration of 10%. Samples were centrifuged using a bench top centrifuge (Eppendorf) at 4000g for 5 min at 4°C. Cell pellets were frozen at -80°C.

2. **Cell lysis.** Yeast cell culture pellets were resuspended in lysis buffer composed of 100 mM triethylamonium bicarbonate (TEAB) containing 8 M Urea and phosphatase/protease inhibitors. Yeast cells were lysed by glass beads beating using a Mini-Beadbeater-24. Samples were shaken in the presence of 0.1 g acid washed glass beads using shock resistant tubes 10 times for 1 min with 1 min rest on ice in-between.

3. **In-solution digestion.** The protein supernatant was separated from insoluble cell debris by centrifugation at 14.000 rpm for 5 min at 10°C using a benchtop centrifuge (Eppendorf). Proteins were reduced by addition of 5 mM Dithiothreitol (DTT) (final concentration) for 1 hour at 37°C using an Eppendorf ThermoMixer. Subsequently, the proteins were alkylated in the dark by addition of 15 mM iodoacetamide for 30 min at room temperature. Protein precipitation was performed by addition of four volumes of ice-cold acetone (-20° C) and proceeded for 1 hour at -20°C. The solution was centrifuged at 14.000 rpm using a bench-top centrifuge at 10°C to collect the cell pellet. The acetone supernatant was discarded. Proteins were then solubilized using 100 mM ammonium bicarbonate (ABC) and proteolytic digestion

was performed by addition of Trypsin (Promega, Madison, WI), to a 1:100 enzyme to protein ratio (v/v), and incubated at 37°C overnight.

4. **Solid phase extraction of the proteolytic digest.** Solid phase extraction of the proteolytic digest was performed with an Oasis HLB 96-well µElution plate (Waters, Milford, USA). The columns were first washed with 750 µL methanol and equilibrated with 1000 µL water. Samples were then loaded and washed twice with 5% MeOH. Peptide fractions were eluted in two 80% MeOH buffers containing 0.01% TFA and 1 mM ABC, separately. Eluates were collected in 1.5 mL LoBind tubes (Eppendorf) and dried using a SpeedVac concentrator for two hours (temperature was set to 50°C for one hour, and to RT for the second hour). Dried peptides were resuspended in 3% acetonitrile (ACN) / 0.01% TFA prior to MS-analysis to give an approximate concentration of 500 ng per µL.

**DETAILED DESCRIPTION PROTOCOL 13**

1. **Sample biomass collection.** Yeast biomass (1 mL) was collected from steady state cultures. The samples were collected in multifold in trichloroacetic acid (TCA) with a final concentration of 10%. Samples were centrifuged using a bench top centrifuge (Eppendorf) at 4000 g for 5 min at 4°C. Cell pellets were frozen at -80°C.

2. **Cell lysis.** Yeast cell culture pellets were resuspended in lysis buffer composed of 100 mM triethylamonium bicarbonate (TEAB) containing 8 M Urea and phosphatase/protease inhibitors. Yeast cells were lysed by glass beads beating using a Mini-Beadbeater-24. Samples were shaken in the presence of 0.1 g acid washed glass beads using shock resistant tubes 10 times for 1 min with 1 min rest on ice in-between.

3. **Filter-aided digestion.** For filter-aided sample preparation (FASP), proteins were loaded to a filter after bead beating, (Merck-Millipore, Microcon 10 kDa). The protein supernatant was separated from insoluble cell debris by centrifugation at 14.000 rpm for 5 min at 10°C using a benchtop centrifuge (Eppendorf). Proteins were reduced on the filter by addition of 5 mM Dithiothreitol (DTT) (final concentration) for 1 hour at 37°C using an Eppendorf ThermoMixer. Subsequently, the proteins were alkylated in the dark by addition of 15 mM iodoacetamide for 30 min at room temperature. After alkylation, proteins were washed two times with TEAB and two times with ABC buffers. Proteolytic digestion was performed by addition of Trypsin (Promega, Madison, WI), to a 1:100 enzyme to protein ratio (v/v), and incubated at 37°C overnight. Finally, peptides were eluted in two fractions from the column using a 200 mM ABC and a 5% ACN/0.1% formic acid (FA) buffer.

4. **Solid phase extraction of the proteolytic digest.** Solid phase extraction of the proteolytic digest was performed with an Oasis HLB 96-well µElution plate (Waters, Milford, USA). The columns were first washed with 750 µL methanol and equilibrated with 1000 µL water. Samples were then loaded and washed twice with 5% MeOH. Peptide fractions were eluted in two 80% MeOH buffers containing 0.01% TFA and 1 mM ABC, separately. Eluates were collected in 1.5 mL LoBind tubes (Eppendorf) and dried using a SpeedVac concentrator for two hours (temperature was set to 50°C for one hour, and to RT for the second hour). Dried peptides were resuspended in 3% acetonitrile (ACN) / 0.01% TFA prior to MS-analysis to give an approximate concentration of 500 ng per µL

**DETAILED DESCRIPTION PROTOCOL 14**

1. **Sample biomass collection.** Yeast biomass (1 mL) was collected from steady state cultures. The samples were collected in multifold in trichloroacetic acid (TCA) with a final concentration of 10%. Samples were centrifuged using a bench top centrifuge (Eppendorf) at 4000 g for 5 min at 4°C. Cell pellets were frozen at -80°C.

2. **Cell lysis.** Yeast cell culture pellets were resuspended in lysis buffer composed of 100 mM triethylamonium bicarbonate (TEAB) containing 1% SDS and phosphatase/protease inhibitors. Yeast cells were lysed by glass beads beating using a Mini-Beadbeater-24. Samples were shaken in the presence of 0.1 g acid washed glass beads using shock resistant tubes 10 times for 1 min with 1 min rest on ice in-between.

3. **Filter-aided digestion.** For filter-aided sample preparation (FASP), proteins were loaded to a filter after bead beating, (Merck-Millipore, Microcon 10 kDa). The protein supernatant was separated from insoluble cell debris by centrifugation at 14.000 rpm for 5 min at 10°C using a benchtop centrifuge (Eppendorf). Proteins were reduced on the filter by addition of 5 mM Dithiothreitol (DTT) (final concentration) for 1 hour at 37°C using an Eppendorf ThermoMixer. Subsequently, the proteins were alkylated in the dark by addition of 15 mM iodoacetamide for 30 min at room temperature. After alkylation, proteins were washed two times with TEAB and two times with ABC buffers. Proteolytic digestion was performed by addition of Trypsin (Promega, Madison, WI), to a 1:100 enzyme to protein ratio (v/v), and incubated at 37°C overnight. Finally, peptides were eluted in two fractions from the column using a 200 mM ABC and a 5% ACN/0.1% formic acid (FA) buffer.

4. **Solid phase extraction of the proteolytic digest.** Solid phase extraction of the proteolytic digest was performed with an Oasis HLB 96-well µElution plate (Waters, Milford, USA). The columns were first washed with 750 µL methanol and equilibrated with 1000 µL water. Samples were then loaded and washed twice with 5% MeOH. Peptide fractions were eluted in two 80% MeOH buffers containing 0.01% TFA and 1 mM ABC, separately. Eluates were collected in 1.5 mL LoBind tubes (Eppendorf) and dried using a SpeedVac concentrator for two hours (temperature was set to 50°C for one hour, and to RT for the second hour). Dried peptides were resuspended in 3% acetonitrile (ACN) / 0.01% TFA prior to MS-analysis to give an approximate concentration of 500 ng per µL.
